# Guided construction of single cell reference for human and mouse lung

**DOI:** 10.1101/2022.05.18.491687

**Authors:** Minzhe Guo, Michael P. Morley, Yixin Wu, Yina Du, Shuyang Zhao, Andrew Wagner, Michal Kouril, Kang Jin, Nathan Gaddis, Joseph A. Kitzmiller, Kathleen Stewart, Maria C. Basil, Susan M. Lin, Yun Ying, Apoorva Babu, Kathryn A. Wikenheiser-Brokamp, Kyu Shik Mun, Anjaparavanda P. Naren, Sara Lin, Geremy Clair, Joshua N. Adkins, Gloria S. Pryhuber, Ravi S. Misra, Bruce J. Aronow, Timothy L. Tickle, Nathan Salomonis, Xin Sun, Edward E. Morrisey, Jeffrey A. Whitsett, Yan Xu, NHLBI LungMAP Consortium

## Abstract

Accurate cell type identification is a key and rate-limiting step in single cell data analysis. Single cell references with comprehensive cell types, reproducible and functional validated cell identities, and common nomenclatures are much needed by the research community to optimize automated cell type annotation and facilitate data integration, sharing, and collaboration. In the present study, we developed a novel computational pipeline to utilize the LungMAP CellCards as a dictionary to consolidate single-cell transcriptomic datasets of 104 human lungs and 17 mouse lung samples and constructed “LungMAP CellRef” and “LungMAP CellRef Seed” for both normal human and mouse lungs. “CellRef Seed” has an equivalent prediction power and produces consistent cell annotation as does “CellRef” but improves computational efficiency and simplifies its utilization for fast automated cell type annotation and online visualization. This atlas set incorporates 48 human and 40 mouse well-defined lung cell types catalogued from diverse anatomic locations and developmental time points. Using independent datasets, we demonstrated the utility of our CellRefs for automated cell type annotation analysis of both normal and disease lungs. User-friendly web interfaces were developed to support easy access and maximal utilization of the LungMAP CellRefs. LungMAP CellRefs are freely available to the pulmonary research community through fast interactive web interfaces to facilitate hypothesis generation, research discovery, and identification of cell type alterations in disease conditions.

## INTRODUCTION

Single cell RNA-seq (scRNA-seq) analysis is being widely applied to biomedical research, enabling the study of complex organs, such as the lung, at unprecedented scale and resolution, and transforming our understanding of organ development and disease^1-4^. Accurate cell type identification is a key and rate-limiting step in the single cell data analysis that usually requires time-consuming processes to optimize computational parameters followed by manual inspections that require domain expertise. With the increasing number of published scRNA-seq datasets and the release of large-scale cell atlases, advanced computational tools^5-7^ have been developed using annotated datasets to predict cell identities in new datasets. Common issues related with the use of a published scRNA-seq data as a reference for supervised classification of user-supplied datasets include the lack of comprehension (missing cell types), inclusion of speculative cell types/states that have not been functionally validated, technology specific-biases in the reference or query, and insufficiently powered to represent the repertoire of common healthy lung cell types. The lack of common cell type nomenclatures and guidelines for single cell transcriptomic studies also creates substantial technical challenges for data integration and comparison. Therefore, single cell references with comprehensive cell types, reproducible and functional validated cell identities, and common nomenclatures are much needed by the research community to optimize automated cell type annotation and facilitate data integration, sharing and collaboration.

A growing number of community-wide efforts have been devoted to the development of common cell type nomenclatures, including cell type ontologies of the Human Cell Atlas^8^ and the common cell type nomenclature for the mammalian brain^9^. Recently, the LungMAP consortium produced a LungMAP CellCards^10^, a rigorous catalogue of lung cells based on a community effort of the consortium that synthesizes current functional and single cell data from human and mouse lungs into a comprehensive and practical cellular census of lung cells. The current version of LungMAP CellCards catalogued 39 major cell types and numerous immune cell subtypes, spanning the principal cellular heterogeneity present in diverse regions of normal lung, including trachea, bronchi, submucosal glands (SMG), and lung parenchyma^10^. These common cell type nomenclature efforts provide a scaffold and guideline for the ongoing development of a comprehensive lung single cell reference for single-cell genomics analysis. In addition to curation, novel computational methods are further needed to utilize these common cell type nomenclatures as guidelines to accurately identify cell types using integrated single cell datasets.

Here, we present a novel approach for cell atlas construction that directs the identification of reference cell populations according to a dictionary of pre-compiled cell type terms and molecular markers derived from CellCards. The pipeline consists of two key steps, first identifying a “seed” population for each cell type which best represents the cell identity in the dictionary, then mapping all cells to the “seeds” based on transcriptomic similarity to construct a complete single cell reference, term “CellRef”. Using this approach, we constructed and released a CellRef consisting of a total of 48 normal human lung cell types, which we named “LungMAP Human Lung CellRef”. Using the same approach, we identified “seed” cells for 40 mouse lung cell types and constructed the “LungMAP Mouse Lung Development CellRef”. We deployed this resource as multiple user-friendly web interfaces to facilitate easy access and maximal use of the LungMAP CellRefs. These interfaces include use of the recently developed Azimuth interface, which enables research investigators to annotate their own scRNA-seq dataset automatically using the LungMAP CellRef, via automated supervised classification, prior user-annotation comparison, and exploration against the CellRef for any scRNA-seq input dataset (up to 100,000 cells). We developed functions to facilitate evaluation of automated cell type annotation results using CellRef marker genes. Using prior published datasets, we demonstrated the utility of LungMAP CellRefs for automated cell type annotation analysis of scRNA-seq data from normal and diseased human lungs. The present guided approach is implemented in R and is applicable for CellRef construction for other organs.

## RESULTS

### Data collection and guided construction for LungMAP single cell reference

The LungMAP CellCards catalogued 39 cell types and their associated marker genes in multiple regions of normal lung, including trachea, bronchi, SMG, and lung parenchyma^10^. To construct a LungMAP human lung CellRef in accordance with the CellCards, we collected 10 large-scale sc/snRNA-seq datasets (8 published and 2 unpublished) from the four regions of human lung (**Figure 1A**): Habermann et al.^11^ (n=10 donors; parenchyma), Reyfman et al.^12^ (n=8 donors; parenchyma), Adams et al.^13^ (n=28 donors; parenchyma), Deprez et al.^14^ (n=9 donors; trachea/bronchi/parenchyma), Travaglini et al.^15^ (n=3 donors; bronchi/parenchyma), Goldfarbmuren et al.^16^ (n=15 donors; trachea), Wang et al.^17^ (n=3, small airway), Melms et al.^3^ (n=7, parenchyma), CCHMC LungMAP cohort (n=5, bronchus SMG) and UPenn LungMAP cohort (n=16, parenchyma). This collection contains data from similar numbers of female and male donors (n=48 and 55, respectively; 1 unannotated) (**Figure 1A; Supplementary Table 1**). The median age of donors was 41 years (interquartile range [IQR], 29 - 61 years; 1 unannotated). Data were generated from three 10x chromium single cell libraries: Single Cell 3’ sequencing kit based on v2/v3 and Single Cell 5’ chemistry. In total, sc/snRNA-seq of 505,256 lung cells from 148 normal human lung samples from 104 donors were used for LungMAP human lung CellRef construction (**Supplementary Table 1**).

**Figure 1.**
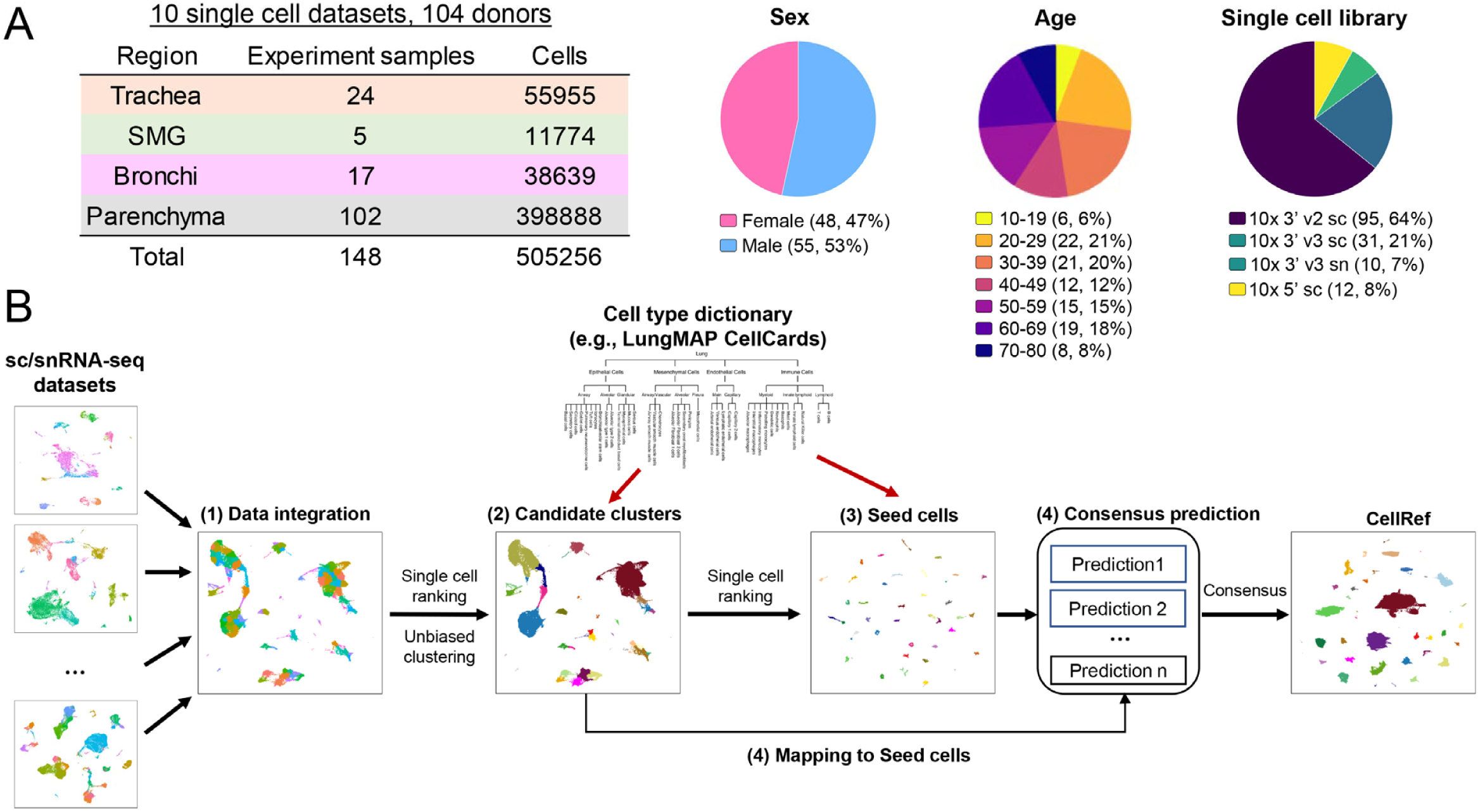
Data collection and the guided single cell reference (CellRef) construction pipeline. (**A**) Characteristics of the collection of single cell/nucleus RNA-seq datasets from normal human lung. (**B**) Schematic workflow for the LungMAP CellRef construction guided by using LungMAP CellCards as a cell type dictionary.

The integration of such a large and complex single cell data collection is extremely challenging due to the huge batch differences associated with both biological (i.e., different donor and different anatomic regions) and technical variations (e.g., sample preparations by different protocols from different research institutions). To perform accurate single cell reference construction, we developed a novel computational pipeline which combines batch correction, unsupervised cell clustering, “single cell ranking”, power analysis, and automated cell type annotation to consolidate single cell datasets and annotate cell identities guided by a pre-defined cell type dictionary (i.e., LungMAP CellCards) (**Figure 1B**). We utilized both positive and negative markers to improve the sensitivity to distinguish cell types sharing similar gene expression patterns and marker genes, for example, lung goblet cells (*MUC5AC+/MUC5B+*) and SMG mucous (*MUC5AC*-*/MUC5B*+). The pipeline consists of four major steps (**Figure 1B, Methods**). First, batch correction. We used the mutual nearest neighbor (MNN) matching method in Monocle 3 as default, and Seurat’s reciprocal principal component analysis (RPCA) based integration^5^ and Harmony^18^ as alternative. Next, “seed” identification (steps 2 and 3 in **Figure 1B**). This is a unique feature of our approach. We aim to identify a core set of cells that best match to the identity of each cell type in the dictionary. We perform unbiased clustering analysis and determine candidate cell clusters for each cell type based on the expression of marker genes. The use of unbiased clustering provides an opportunity to discover new cell types that are not yet defined in the dictionary. To identify the best “seed” cells, we developed a “single cell ranking” method that first ranks cells based on expression of each cell specific marker gene in the dictionary and then aggregates the rankings of all markers for a given cell type to identify “seed” cells for the cell type. We performed a power analysis to determine the minimum number of “seed” cells required. The last step is automated cell type annotation. We applied multiple cell type annotation methods (e.g., Seurat’s label transfer and SingleR) to map all other cells to the “seed” cells and predict their cell types. Cells that have consistent cell type predictions in all methods will be included in the CellRef. The last step can be repeated to include newly collected datasets into the CellRef by mapping them to the “seed” cells. We implemented this cell-type-dictionary guided CellRef construction pipeline in R and hosted its development and documentation in github: https://github.com/xu-lab/CellRef.

### The LungMAP Human Lung CellRef

Using this guided approach and a cell type dictionary derived from LungMAP CellCards (**Supplementary Table 2**), we identified 8,080 “seed” cells representing 48 normal human lung cell types, termed “**LungMAP Human Lung CellRef Seed**” (**Figure 2A**). Next, we mapped all other cells in our collection to the “seed” cells and predicted cell type annotations using two independent methods, “Seurat Label Transfer” and “SingleR”. Cells with consistent cell type annotations were combined with the “seeds” to form the “**LungMAP Human Lung CellRef**” (347,970 cells) (**Figure 2B, Supplementary Figures 1-2, Methods**).

**Figure 2.**
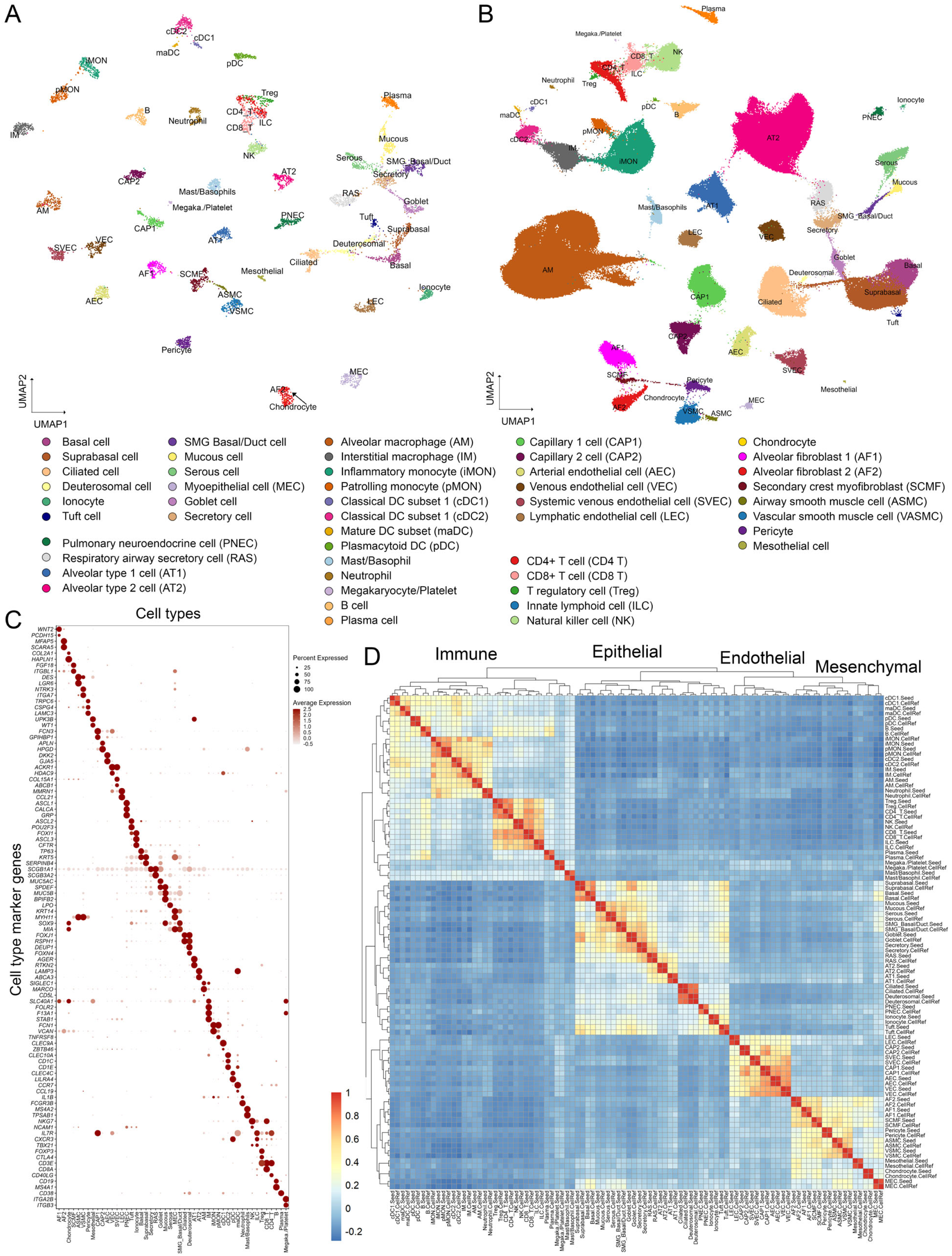
The construction of “LungMAP Human Lung CellRef”. (**A**) Uniform manifold approximation and projection (UMAP) visualization of “seed” cells representing 48 lung cell types of normal human lung, termed “LungMAP Human Lung CellRef Seed”. Cells were colored by their predicted seed identities. (**B**) UMAP visualization of the complete single cell reference for normal human lung, denoted as “LungMAP Human Lung CellRef”, which contains 347,970 cells from 104 donors and defines 48 cell types in normal human lung. Cells were colored by their predicted identities. (**C**) Validation of the “seed” cell identity using the expression of cell type selective marker genes derived from LungMAP CellCards. (**D**) Reconstruction of cell lineage relationships using hierarchical clustering analysis of cell type pseudo-bulk gene expression profiles. Color represents Pearson’s correlation of pseudo-bulk expression profiles. Labels ending with “.Seed” represent pseudo-bulk profiles created by averaging gene expression in the cells of each cell type in the “LungMAP Human Lung CellRef Seed”, while labels ending with “.CellRef” represent pseudo-bulk profiles created using gene expression of each cell type in the “LungMAP Human Lung CellRef”.

The CellRef includes the following CellCards cells: 12 epithelial (AT1, AT2, basal, ciliated, goblet, myoepithelial [MEC], mucous, PNEC, secretory, serous, Tuft cells, and ionocytes); 5 endothelial (arterial, venous, lymphatic endothelial and capillary 1 and 2 cells), 8 mesenchymal (alveolar fibroblast 1 and 2 [AF1, AF2], airway and vascular smooth muscle cells [ASMC, VSMC], mesothelial, chondrocytes, pericytes, and myofibroblasts [SCMF]), and 16 immune cell types (alveolar and interstitial macrophage [AM, IM], inflammatory and patrolling monocytes [iMON, pMON], mast/basophils, neutrophils, B, plasma, NK, ILC, cDC1, cDC2, pDC, CD8+ T, CD4+ T, and T regulatory [Treg] cells). In addition to the known lung cell types, we extended the dictionary to incorporate 7 cell types that are not yet in the CellCards but have marker genes reported in recent scRNA-seq studies and selectively expressed in our unbiasedly identified cell clusters, including deuterosomal cells^14^ (*DEUP1, FOXN4, CDC20B*), suprabasal cells^14^ (*SERPINB4, KRT19, NOTCH3*), systemic venous endothelial cells^19^ (SVEC; marker genes: *COL15A1, ABCB1, ACKR1*), mature dendritic cell subset^20^ (maDC; marker genes: *CCR7, CCL19, LAD1*), megakaryocyte/platelets^15,21^ (*ITGA2B, ITGB3*), SMG duct cells^22^ (*MIA, ALDH1A3, RARRES1*), and respiratory airway secretory cells^4^ (RAS; marker genes: *SCGB3A2, KLK11, SOX4*). We combined SMG basal and SMG duct cells into one mixed type, “SMG Basal/Duct cell”, since their marker genes were co-expressed in the same cell cluster in our data integration. Similarly, we combined mast and basophil cells into a mixed “Mast/Basophil” cell type. We performed uniform manifold approximation and projection for dimension reduction (UMAP) analysis on the “LungMAP Human Lung CellRef”. All cells, from trachea to alveoli, were projected into a common UMAP space and showed clear separations by the predicted cell identities (**Figure 2B**).

To evaluate cell identities in the “LungMAP Human Lung CellRef Seed”, we preformed the following validation analyses. Cell type marker genes were found to be selectively expressed in their corresponding “seed” cells, the majority having high cell type specific expression frequencies, suggesting that the cell identities of the “seed” cells were consistent with the cell type dictionary (**Figure 2C**). To further validate the identities of the “seed” cells, we created pseudo-bulk gene expression profiles for each cell type by averaging gene expression in its “seed” cells, measured their correlations, and performed hierarchical clustering analysis, demonstrating that cell types were first unbiasedly clustered by their major cell lineages and then by sub-lineages (**Figure 2D)**. The pseudo-bulk profile of SMG myoepithelial cells (MEC) co-clustered with mesenchymal cells and positively correlated with both the profiles of SMG Basal/Duct cells and smooth muscle cells, consistent with their complex cell nature. UMAP analysis showed that the “seed” cells formed dense cell clusters and clearly distinguished all cell types except closely related T cell subtypes (i.e., Treg and ILC are clustered with CD8/4 T cells), supporting distinct transcriptomic patterns of cell types in the “LungMAP Human Lung CellRef Seed” and a high similarity of the “seed” cells representing each cell type (**Figure 2A**). In summary, using our guided approach, we developed the “LungMAP Human Lung CellRef Seed”, a collection of “seed” cells for 48 normal lung cell types with cell identities in accordance with a cell type dictionary derived from the LungMAP CellCards.

To validate the similarity of cell identities in the “seed” and “CellRef’, we created pseudo-bulk profiles for the cell types in the “LungMAP Human Lung CellRef”, combined them with the pseudo-bulk profiles generated using the “seed” cells, measured correlations among all pseudo-bulk profiles, and performed unbiased hierarchical clustering analysis. Like the “seed” cells, the pseudo-bulk profiles of the cell types in the “LungMAP Human Lung CellRef” were also first clustered by their major cell lineages and then by sub-lineages. Moreover, each of them was well correlated with the pseudo-bulk profile of the same cell type created using the “seed” cells (**Figure 2D**). Taken together, these results validated the identities of cell types in our constructed “LungMAP Human Lung CellRef”.

### The LungMAP Mouse Lung Development CellRef

Using the above approach, we constructed a cell type dictionary based on the LungMAP CellCards to define cell types in mouse lung during perinatal development, identified “seed” cells for each cell type (termed “LungMAP Mouse Lung Development CellRef Seed”), and constructed a CellRef for mouse lung development (denoted as “LungMAP Mouse Lung Development CellRef”) using Drop-seq of mouse lungs (n=95,658, 17 experimental samples, 8 time points) from embryonic day 16.5 to postnatal day 28 (**Figure 3, Supplementary Figure 3, Supplementary Tables 3-4**). Because of the time course design, the mouse lung CellRef included more developmental progenitor cells and transitional cell states than the LungMAP Human Lung CellRef, including, *Sox9*+/*Id2*+ distal epithelial progenitor cell^23,24^, an “AT1/AT2” cell population^21,25^ expressing both AT1 (*Ager, Hopx*) and AT2 (*Lamp3, Sftpc, Abca3*) cell markers in conjunction with *Cldn4, Krt19*, and *Krt8* (signature genes of recently reported PATS^26^, DATP^27^, or ADI^28^ cells), *Foxf1*+/*Kit*+ endothelial progenitor cells (EPC) ^29^, and proliferative mesenchymal progenitor (PMP) cells^30,31^ (**Figure 3**). In total, 40 mouse lung cell types have been identified with the guidance of the mouse lung cell type dictionary (**Figure 3, Supplementary Table 4**). Cell identities were validated using expression of marker genes, UMAP visualization of cell types, pseudo-bulk expression and hierarchical clustering analysis based cell lineage reconstruction, and cell type specific signature gene identification (**Figure 3**). The construction of this LungMAP mouse lung CellRef in parallel with the human lung CellRef will enable cross comparisons for better understanding of how the cell types in mouse lung relate to the human lung and how data from mouse studies in the literature relate to human disease.

**Figure 3.**
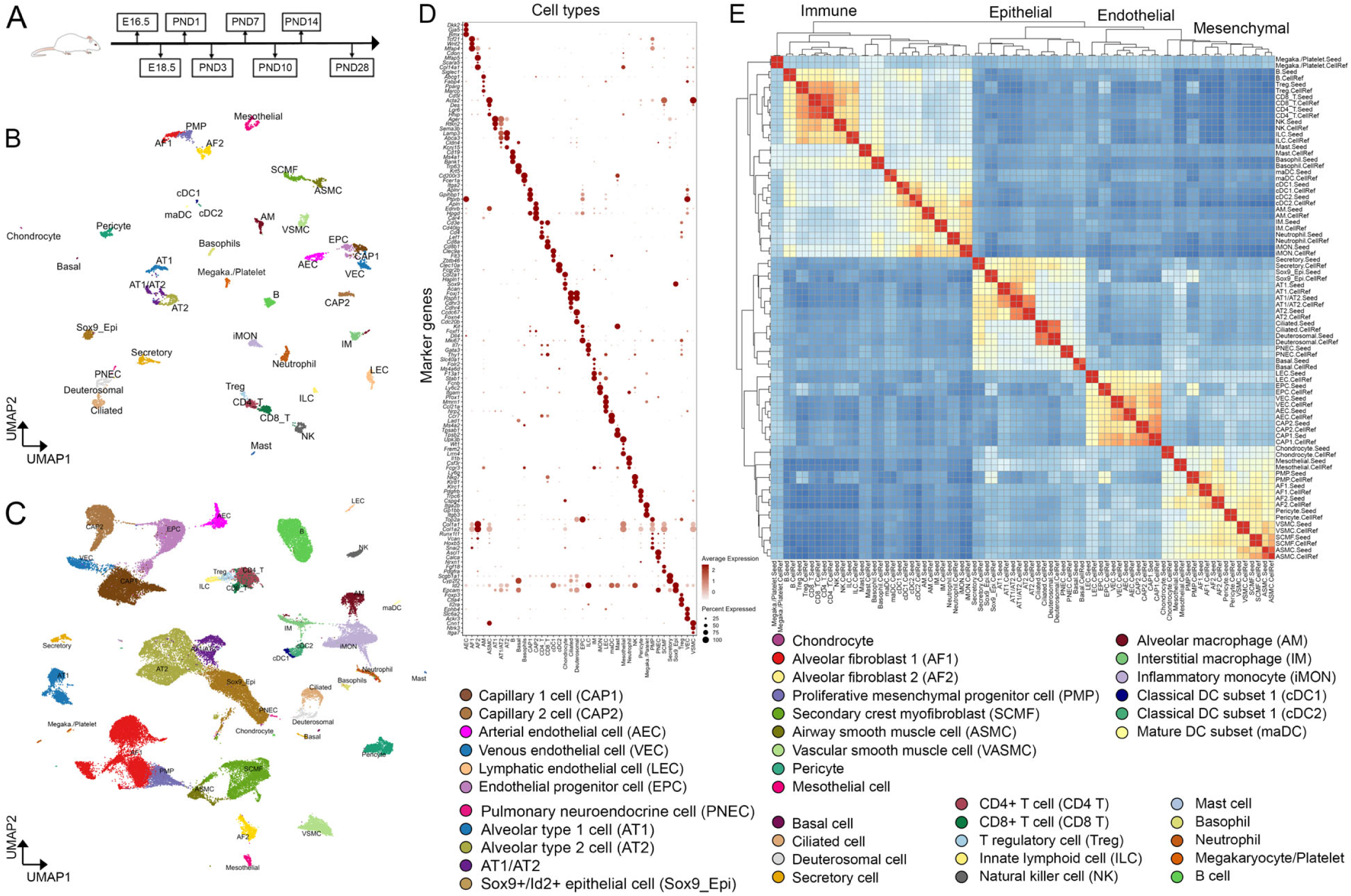
The construction of “LungMAP Mouse Lung Development CellRef”. (**A**) The developmental time points of mouse lung single cell transcriptome data used for the guided CellRef construction. (**B**) Uniform manifold approximation and projection (UMAP) visualization of the “seed” cells representing 40 cell types of the developing mouse lung, termed “LungMAP Mouse Lung Development CellRef Seed”. Cells were colored by predicted seed identities. (**C**) UMAP visualization of CellRef for normal mouse lung development, named “LungMAP Mouse Lung Development CellRef”. Cells were colored by their predicted identities. (**D**) Validation of cell type identities in “LungMAP Mouse Lung Development CellRef Seed” using expression of cell type selective marker genes derived from the LungMAP CellCards. (**E**) Lineage relationships among mouse lung cell types were reconstructed using hierarchical clustering analysis using pseudo-bulk gene expression profiles.

### Interactive web-tools for search and display of the LungMAP CellRefs

To facilitate data sharing and broad use of the resource, we developed several user-friendly web portals to host the LungMAP single cell references online, including the LGEA LungMAP CellRef page (https://research.cchmc.org/pbge/lunggens/CellRef/LungMapCellRef.html) and scViewer-Lite (http://devapp.lungmap.net/app/seuratviewer-lite, an R Shiny based application). These tools provide highly interactive search and visualization functionalities for users to explore cell type and gene expression patterns provided by the LungMAP CellRefs (**Figure 4, Supplementary Figure 4**). The LGEA CellRef page enables users to perform “Gene Expression Query”, “Cell Type Query”, and “Cell Signature Query”. The “Gene Expression Query” enables users to input any gene of interest to visualize the expression patterns and associated statistics in UMAP, Box, Notched Box, Beeswarm, Scatter plot, and bi-directional bar charts (**Figure 4A-B**). The “Cell Type Query” enables users to select any one of the pre-defined cell types and obtain cell-type information collected by LGEA^32^ including cell selective marker genes, transcription factors, and surface markers, ligands and receptors) as well as a link to the LungMAP CellCards^29^ (**Figure 4C**). The “Cell Signature Query” function provides cell type selective signature genes identified using the LungMAP CellRefs, along with interactive tables and bar graph that enables users to search differential expression statistics and compare the mean gene expression across all cell types (**Figure 4D**). scViewer-lite is a R shiny based app that allows for comparative viewing of gene expression and/or other meta data overlapped on dimension reduction plots and violin plots. Users can also select and highlight cells of interest (**Supplementary Figure 4**). In addition to these two newly developed web interfaces, LungMAP CellRefs can be interactively explored in LungMAP web portal using ShinyCell^33^ based web interfaces (https://lungmap.net/cell-cards/, “scRNA-seq” tab).

**Figure 4.**
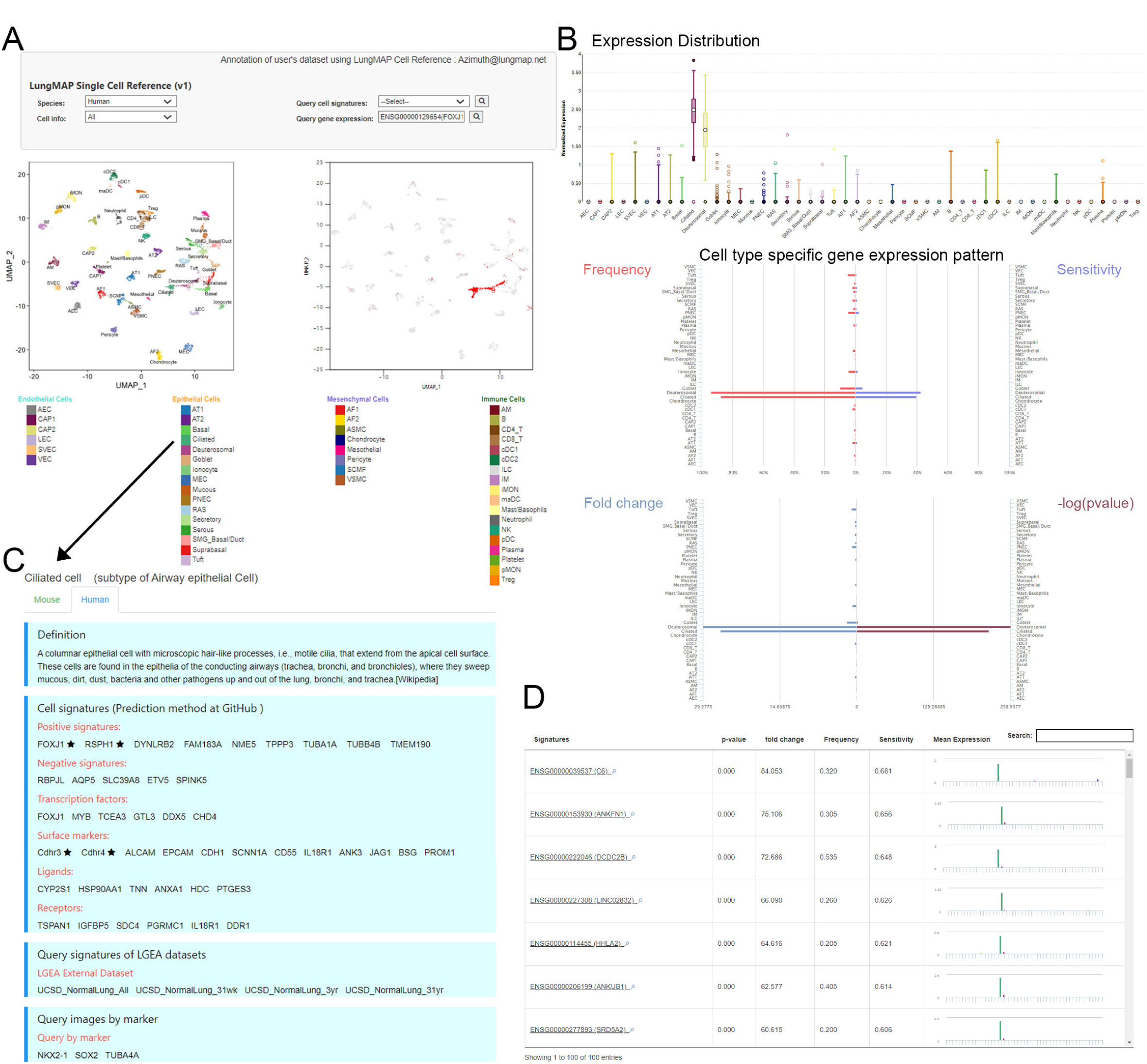
Online interactive exploration of LungMAP CellRef Seed using Lung Gene Expression Analysis (LGEA) web portal. The “LungMAP Human Lung CellRef Seed” was comprised of 8,080 “seed” cells representing 48 normal lung cell types. (**A**) The “Gene Expression Query” interface allows users to input a gene of interest (top) and visualize of the expression of the queried gene in UMAP embeddings of cells (bottom), Colors represent the putative cell identities (bottom left) or the expression of the input gene (bottom right). (**B**) Visualization of the gene expression pattern (top: expression distribution; middle: expression frequency and sensitivity; bottom: fold change and p-value of differential expression) across all cell types in the CellRef Seed. (**C**) LGEA hosts comprehensive cell information related with the query cell type. (**D**) “Cell Signature Query” function retrieves signature gene expression statistics of a given cell type and bar-plot visualization of signature genes expression across all cell types in the “LungMAP Human Lung CellRef Seed”. In (A) and (B), *FOXJ1* expression was shown as example. In (C) and (D), “Ciliated cells” were used as example.

### Automated cell type annotation using the LungMAP CellRefs

We developed LGEA and scViewer-lite for users to explore expression patterns of normal lung cells and genes of interest without the need for computational coding. Another powerful use of the LungMAP CellRefs is to use them for automated cell type annotation of users’ own single cell datasets to facilitate analysis and standardization of cell type prediction and annotations. To achieve this goal, we built our LungMAP “seed” cells and CellRefs into R objects in accordance with Seurat reference mapping pipeline^5,34^. Azimuth (https://satijalab.org/azimuth/) instances were established at LungMAP.net (https://lungmap.net/cell-cards/, “Azimuth” tab) to enable users to upload their own datasets (up to 100,000 cells) for online automated cell type annotation using our CellRefs (**Figure 5A**) and exploration of any gene features on the projected UMAP or in Violin plots. Additionally, to facilitate evaluation of automated cell type annotation results, we developed functions in our R pipeline to visualize the expression of CellRef markers across all predicted cell types, identify cell type signature genes and their associated functional annotations, and compile all visualization and evaluation results into a single evaluation report using R markdown (**Figure 5A**).

**Figure 5.**
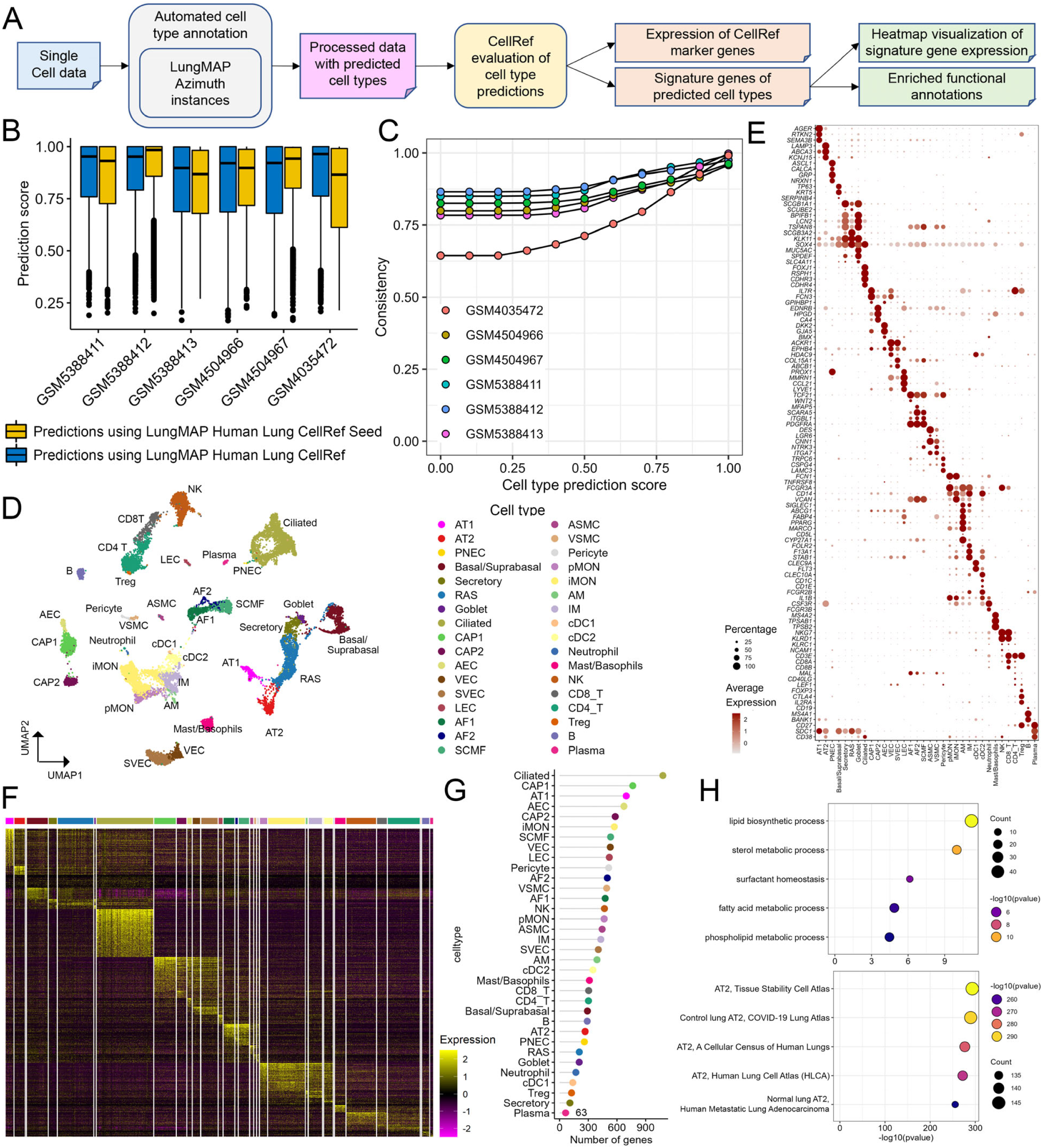
Cell type annotation and evaluation using the LungMAP Human Lung CellRef. (**A**) Schematic workflow of the automated cell type annotation and evaluation pipeline. (**B**) Distributions of cell type prediction scores in each test dataset. Prediction scores using “LungMAP Human Lung CellRef Seed” (yellow bars) are comparable to those using the complete “LungMAP Human Lung CellRef” (blue bars). (**C**) Consistent cell type predictions using “LungMAP Human Lung CellRef Seed” and using “LungMAP Human Lung CellRef” in the six test datasets. Consistency percentages (y axis) were calculated for cells in each test dataset (color). X axis: prediction scores. (**D-H**) Evaluation of automated cell type annotations for three of our test datasets (GSM5388411/12/13, three scRNA-seq of normal human lungs). Evaluation of the other three testing datasets were shown in Supplementary Figure 5. (**D**) UMAP visualization of cells with prediction scores >=0.6 and predicted annotations with at least 5 cells. Cells were colored by automated cell type annotations using the “LungMAP Human Lung CellRef Seed” as reference. Data from different donors were integrated using Seurat’s RPCA pipeline. (**E**) Evaluation of cell type annotations using expression of marker genes derived from LungMAP CellCards. (**F**) Heatmap visualization of expression of cell type specific differentially expressed genes (DEGs). **(G)** The number of DEGs for each predicted cell type annotation. (**H**) Significantly enriched functional annotations using DEGs of the predicted AT2 cells: most enriched “Gene Ontology Biological Processes” (top) and “ToppCell Gene Sets” (bottom). Functional enrichment analysis was performed using ToppGene software suite.

### Use cases driven evaluation of LungMAP single cell references for automated cell type annotation

*Annotation of scRNA-seq of normal human lung*. Six published scRNA-seq datasets of normal human lung samples were collected to independently evaluate the accuracy of the automated cell type annotation using the LungMAP CellRefs. The datasets were from normal human lung samples of 2 months to 45 years of age and were generated using 10X chromium 3’ (GSM5388411/12/13^35^ and GSM4504966/67^36^; aligned to hg38 reference genome) and 5’ (GSM4035472^1^; aligned to hg19 reference genome) platforms. For each test dataset, we used both the “LungMAP Human Lung CellRef Seed” and the “LungMAP Human Lung CellRef” to predict cell type annotations using Azimuth’s mapping algorithm^5^. A prediction score (between 0 and 1) was assigned to each cell in the test dataset based on its transcriptomic similarity to cells in the reference. With the minimal prediction score threshold set as >= 0.6, on average, 85.13% cells in the test dataset can be confidently annotated using the “LungMAP Human Lung CellRef” in comparison with 85.10% using the “LungMAP Human Lung CellRef Seed”, suggesting that similar numbers of cells can be confidently predicted when using both the complete “CellRef” and the “CellRef Seed” (**Figure 5B**). In addition, the predicted cell type annotations were highly consistent in all test datasets using both the “CellRef Seed” and the “CellRef” (**Figure 5C**). Predictions using the “LungMAP Human Lung CellRef Seed” were computationally efficient, taking approximately 1 minute to annotate a 10x chromium scRNA-seq of 4000-8000 cells.

We applied the validation functions in our pipeline (**Figure 5A, Methods**) to evaluate the accuracy of cell type predictions for the five test datasets. As shown in **Figure 5D-H** and **Supplementary Figure 5**, predicted cell types were well separated and formed clusters. Cell-type-specific marker genes from CellCards were selectively expressed in each predicted cell type, supporting the concordance of the cell identities (**Figure 5E, Supplementary Figure 5**). Cell type specific signature genes were identified using widely accepted criteria (adjusted p value of Wilcoxon rank sum test <0.1, expression frequency >=20%, and fold change >=1.5) (**Figure 5F-G**). Functional enrichment analysis of cell type signature genes was used to further validate the predicted cell identities. For example, predicted AT2 cells were functionally enriched in “surfactant homeostasis” and “lipid/phospholipid/fatty acid metabolic processes”. ToppCell (ttps://toppcell.cchmc.org/) analysis showed that the predicted signature genes were consistent with genes selectively expressed in normal AT2 cell identified in independent single cell studies of human lung^37,38^ (**Figure 5H**). In summary, these evaluations demonstrated the general applicability of our constructed single cell references for annotating cell types in scRNA-seq datasets of normal human lung.

*Application to scRNA-seq of human lung diseases*. We previously performed single cell transcriptomic analyses of lung samples from patients with lymphangioleiomyomatosis^1^ and identified a unique population of cells termed LAM^CORE^ that were readily distinguished from endogenous lung cell types and shared closest transcriptomic similarity to uterine myocytes in both normal and LAM uteri^1^. In the present work, we re-aligned the data to the hg38 reference genome and performed automated cell type annotation using the “LungMAP Human Lung CellRef Seed”. Using a criterion of prediction score >=0.8, we predicted 31 cell types from the two LAM lungs (**Figure 6A-C**). Cell type predictions were largely consistent with the original clustering-based annotations^1^. Using the “LungMAP Human Lung CellRef Seed”, however, more cell sub-types can be distinguished. Importantly, the previously identified LAM^CORE^ cells (73 cells) had low prediction scores below the cutoff line (**Figure 6D**), supporting the notion that this LAM-associated cell population was not similar to normal lung cell types in the present LungMAP CellRef. Therefore, our constructed LungMAP CellRefs can be used to assist in analysis of lung disease data and identify potential disease related cell clusters.

**Figure 6.**
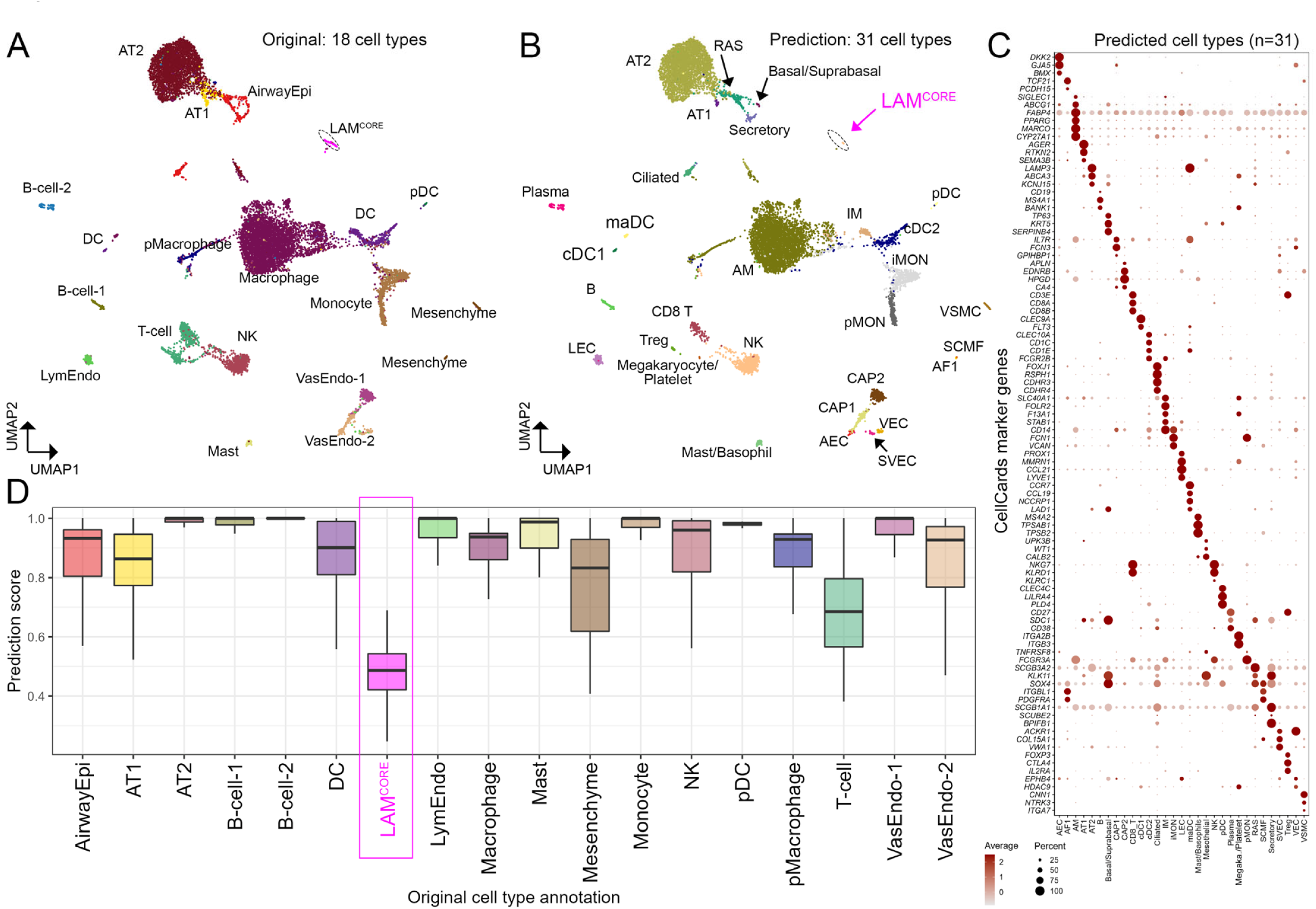
Application of LungMAP Human Lung CellRef to a disease dataset: scRNA-seq of human lungs with lymphangioleiomyomatosis (LAM). (**A**) UMAP visualization of a published scRNA-seq of human LAM lungs. Cell colors represent cell identities predicted in the original publication. A unique disease related cell population, termed as LAM^CORE^ cells (magenta cell cluster) was identified. (**B**) UMAP visualization of cell type annotations using the “LungMAP Human Lung CellRef Seed” as reference. Cells with prediction score >=0.8 were shown. (**C**) Evaluation of cell type annotations using expression of CellRef marker genes. (**D**) Distributions of the cell type prediction scores in the 18 cell types identified in the original study^1^. The disease associated LAM^CORE^ cells had prediction scores below the cutoff line when mapping to LungMAP Human Lung CellRef, suggesting low transcriptomic similarities of LAM^CORE^ cell to normal lung cells in LungMAP Human Lung CellRef.

## DISCUSSION

In the present study, we developed a computational approach to integrate large scale and heterogeneous sc/snRNA-seq datasets and constructed a single cell reference, termed “CellRef”, in accordance with a well-defined cell type dictionary. Using the pipeline and the recently published LungMAP CellCards^10^, we constructed LungMAP CellRefs for normal human lung and developmental mouse lung, termed “LungMAP Human Lung CellRef” and “LungMAP Mouse Lung Development CellRef”, respectively. User-friendly web interfaces were developed to facilitate access, visualization, and use of the LungMAP CellRefs. For advanced users who are interested in annotating their own datasets using the LungMAP CellRefs, we established “Azimuth” instances to support online automated cell type annotations of users’ own scRNA-seq. Evaluation functions were developed in our pipeline to perform fast and comprehensive evaluation of the predicted cell type annotations.

The “LungMAP Human Lung CellRef” contains a total of 347,970 cells and 48 well defined lung cell types, covering major cellular heterogeneity in the four regions: trachea, bronchi, SMG, and lung parenchyma. The “CellRef” identified cell types mapped to the cell type nomenclature in the LungMAP CellCards^10^. In addition, based on unbiased clustering analysis, we identified cell types that are not yet included in the CellCards but reported in recent scRNA-seq analyses, including deuterosomal cells^14^, suprabasal cells^14^, systemic venous endothelial cells^19^, mature dendritic cell subset^20^, SMG duct cells^22^, respiratory airway secretory cells (RAS, a recently identified multipotent secretory cell population in respiratory bronchioles)^4^, and megakaryocyte/platelets^15,21^. During the “CellRef” construction, we discovered cell clusters selectively expressing marker genes of those new cell types, and thus we have included those cell types into the “LungMAP Human Lung CellRef”. We will continue to incorporate more cell types in accordance with new findings from single cell and/or functional studies.

Two earlier versions of human lung references^15,39^ have been published or are in preprint. We compared and incorporated the first reference into our CellRef construction. We carefully compared all annotated cell types in the recently released integrated version of Human Lung Cell Atlas (HLCA) with our CellRef based on the HLCA marker gene expression. Although not all cell type names are identical, the majority of the HLCA annotated cells align well with a clearly defined cell type in our CellRef. Only two fibroblast sub types in HLCA do not align with a clearly defined cell type of CellRef. In addition, each reference identified several unique lung cell types or states (i.e., don’t align to any given cell cluster in the other reference). Further cross-team discussions and comparison is needed to reach consensus on cell identity and common nomenclature, with the ultimate goal of generating a consensus blueprint of normal human lung cell reference for the research community. The present LungMAP CellRefs has several unique features. 1) We developed a new computational pipeline and a guided approach to construct and evaluate the reference which can be reused for future updates of LungMAP CellRef or references of other organs. 2) The LungMAP CellRef identified cell types in accordance with the LungMAP CellCards^10^, a rigorous catalogue of lung cells validated by both single cell and functional studies. 3) During CellRef construction, we identified the best “seed” populations for each cell type (CellRef Seeds), which was not only used to construct the complete “CellRef” but can be independently used for automated cell type annotation and online visualization with improved computational efficiency and hardware requirements. 4) We constructed LungMAP CellRef for both human and mouse, the two most commonly used species, and provide options for users to use either scRNA-seq or snRNA-seq based CellRef separately based on the input sequences type to achieve better performance on cell type annotation. 5) Web portals have been developed by our LungMAP research centers and LungMAP data coordination center to facilitate the resource sharing and maximal use of the constructed references by the research community.

There are several limitations of the current version of LungMAP CellRef and CellRef Seed. Based on the current pipeline and data collection, the present CellRefs do not yet clearly define immune sub-populations and transitional cell states. Antibody base approaches (e.g., CITE-seq or flow cytometry) may help to improve resolution of immune cell sub-populations or cell states^40^. Future lineage/compartment specific reference constructions will be useful in providing enhanced resolutions and granularities at sub-cell type levels or cell transitional states. The current data collections do not have sufficient statistical power for precise annotation of certain rare lung cell types, e.g., SMG duct cells. Region specific Laser Capture Microdissection (LCM) and cell sorting will be useful in identifying and capturing rare lung cell types and their RNA expression patterns. In summary, we developed a novel computational pipeline utilizing a cell type dictionary to consolidate single cell transcriptomic datasets and constructed “LungMAP CellRef” and “LungMAP CellRef Seed” for normal human and mouse lung. “CellRef Seed” has an equivalent prediction power and produces consistent cell annotation as does “CellRef”, but with significantly improved computational efficiency and hardware requirements facilitating utilization for automated cell type annotation and online visualization, addressing a significant computational challenge for single cell reference applications. Using independent datasets, we demonstrated the utility of the CellRefs for automated cell type annotations of normal lung and for potential identification of disease-related cells based on their deviation from normal pulmonary cells. Our CellRefs, along with the analytic and web-based tools, are freely and widely available to the pulmonary research community to facilitate hypothesis generation, research discovery, and identification of cell type alterations in disease conditions.

## METHODS

### Collection and pre-processing of single cell/single nucleus RNA-seq of human lung

We collected eight published and two unpublished sc/snRNA-seq datasets of human lung for LungMAP human lung single cell reference construction. For the published datasets, unique molecular identifier (UMI) count matrix of gene expression in single cells were downloaded from Gene Expression Omnibus (GEO), European Genome-phenome Archive (EGA), or Synapse.org using the following accession numbers: GSE122960^12^, GSE135893^11^, GSE134174^16^, GSE136832^2^ GSE161382^13^, EGAS00001004082^14^, GSE171524^3^, syn21041850^15^. For all datasets, hg38-alignment based data from normal/control lung samples were used.

For the unpublished UPenn LungMAP cohort, samples of normal de-identified human lungs (n=16) from donors who were not matched for lung transplant were obtained as described previously^4^. scRNA-seq experiments (10x Single Cell 3’ v2 and v3 chemistry) were performed as described in Basil et al., 2022^4^. Sequencing read alignment to the hg38 human genome and UMI matrix generation were performed for each sample using 10x Cell Ranger v3 software.

For the unpublished CCHMC LungMAP cohort, microdissection was performed to isolate submucosal gland (SMG) samples from five de-identified normal human lungs for scRNA-seq experiments using 10x Single Cell 3’ v3 sequencing kit. Sequencing read alignment to the hg38 human genome and UMI based gene expression matrix generation were performed for each sample using 10x Cell Ranger v5.

*Data preprocessing*. For published datasets with original cell type annotations, we included cells selected in the original analyses. For published datasets without original cell type annotations (Reyfman et al., 2019^12^) and unpublished datasets (UPenn LungMAP cohort and CCHMC LungMAP cohort), the following quality control (QC) criteria were applied to cell prefiltering, including 500-7,500 expressed genes, less than 25% of UMIs mapped to mitochondrial genes, and less than 50,000 total UMIs. For scRNA-seq data from “Donor29” in the CCHMC LungMAP cohort, we used 1,500-7,500 as the criterion for the “number of expressed genes” based on its unique cell distributions. After pre-filtering, scrublet^41^ was performed to identify and remove potential doublet cells from each data sample. In total, 505,256 cells from 148 lung tissue samples from 104 donors were used as input for our guided pipeline to construct the single cell reference of normal human lung.

### Mice and Drop-seq of mouse lung development

Animal protocols were approved by the Institutional Animal Care and Use Committee in accordance with NIH guidelines. C57BL6/J mice (Jackson Laboratories), female, embryonic days (E) 16.5, 18.5 to postnatal days (PND) 1, 3, 7, 10, 14, 28, were used for single cell RNA-seq experiments using Drop-seq^42^. All mice were time mated. The presence of a vaginal plug was defined as E0.5. PND1 was defined as 24 ± 6 h after birth. Cell isolation and Drop-seq experiments on mouse lungs were described in Guo et al. 2019^21^. The alignment of paired-end sequence reads to mouse genome (mm10) and the generation of digital expression matrix were processed using Drop-seq tools (https://github.com/broadinstitute/Drop-seq/, version 2.3.0) with default parameters. The expression matrix was generated by counting the number of unique molecular identifiers (UMIs) per gene per cell. In total, single cell gene expression in 17 lung samples from eight time points of mouse lung development were generated. For each data sample, the following pre-processing steps were performed. EmptyDrops^43^ was used to identify cell barcodes with expression profiles significantly deviated from the profiles of empty droplets in each data sample with the parameters: lower=100, FDR<0.01. Filters were then applied to keep cells with 400–7,500 genes, less than 40,000 UMIs, and less than 10% UMIs mapped to mitochondrial genes. Potential doublet cells in each sample were predicted and removed using Scrublet^41^. Ambient background RNAs were cleaned from gene expression in each cell using SoupX^44^ using contamination fractions automatically estimated from data.

### Guided construction of single cell reference

Our guided single cell reference (CellRef) construction workflow consists of four major steps: data integration, candidate cell cluster identification, seed cell identification, and consensus prediction for CellRef. We compiled a cell type dictionary containing a list of cell types and associated marker genes, including positive (selectively expressed in the cell type) and negative (no expression in the cell type) markers. We required at least two positive markers for each defined cell type to be included in our CellRef construction.

i. *Data integration*. Multiple algorithms have been integrated into our R workflow, including mutual nearest neighbor (MNN) matching^45^, reciprocal principal component analysis (RPCA) in Seurat^34^, and Harmony^18^. By default, we use the “align_cds” function in Monocle 3 to perform MNN matching based data integration and batch correction. Before integration, we merge data from all datasets into a single gene expression matrix, use it to construct a Monocle 3 “cell_data_set” object, and use the “preprocess_cds” function in Monocle 3 to normalize data to address read depth differences, regress out cell cycle effects and mitochondrial percentage differences, and calculate principal components representing major variances in the data.
ii. *Candidate cell cluster identification*. Using the integrated data, we identify candidate cell clusters for each cell type listed in the dictionary using a combination of unbiased clustering algorithm and marker based “single cell ranking”. We perform unbiased clustering analysis to group cells into distinct cell clusters based on transcriptomic similarity. By default, we perform clustering using the Leiden algorithm^46^ implemented in the “cluster_cells” function in Monocle 3. Followed by the unbiased clustering analysis, we perform a “single cell ranking” for each cell type *i* listed in the dictionary. Let *P*_*i*_ be the set of positive marker genes of cell type *i*. For each marker gene *x* ∈ *P*_*i*_, we identify *Z*_*xi*_, a set of cells with positive (>0) zscore-scale expression of *x*, and generate *R*_*xi*_, a ranking of cells in *Z*_*xi*_ in the descending order based on zscore-scaled expression of *x*. We then aggregate all rankings {*R*_*xi*_|*x* ∈ *P*_*i*_} into a single global ranking of cells, denoted as *R*_*i*_, for the cell type *i*, aiming to identify cells that are ranked highly by multiple cell type marker genes. The aggregation was performed using an order-statistics-based robust rank aggregation algorithm^47^, which assigns a score to each cell in *R*_*i*_ to represent significance of the cell that is ranked consistently better than expected under a null hypothesis derived from {*R*_*xi*_|*x* ∈ *P*_*i*_}. Cells passing selection criteria were used as candidates for cell type mapping. Using the clustering and single cell ranking results, we determine candidate cell clusters for each cell type *i* as follows. Let *𝜑*_*i*_ be the set of cells passed selection criteria (by default, score <0.1) in *R*_*i*_ and *∑* be the cell clusters that we obtained from the unbiased clustering analysis. We calculate the precision and recall values for each cluster *𝜎*_*j*_ ∈ ∑ as follows: *precision*(*i,j*) = l*𝜑*_*i*_ *∩ 𝜎*_*j*_|/|*𝜎*_*j*_ |, *recall*(*i, j*) = |*𝜑*_*i*_ *∩ 𝜎*_*j*_|/|*𝜑*_*j*_|, where |*𝜑*_*j*_| and |*𝜎*_*j*_l denote the number of cells in *𝜑*_*j*_ and *𝜎*_*j*_, respectively, and |*𝜑*_*i*_ *∩ 𝜎*_*j*_l denotes the number of cells in both *𝜎*_*j*_ and *𝜑*_*j*_. The candidate cell clusters for cell type *i* is determined as *A*_*i*_ = {*𝜎*_*j*_ ∈ ∑|*precision*(*i, j*) ≥. *F, recall*(*i, j*) ≥. *S, F* ∈ [0,1], *S* ∈ [0,1]}. A QC inspection of the candidate cell clusters is recommended to ensure the accuracy for the CellRef construction. In summary, in step 2, we use unbiased clustering in conjugation with marker based single cell ranking to select most relevant cell groups as candidates. The use of unbiased clustering before seed cell identification can also provide an opportunity to discover new cell types that have not yet been defined in the dictionary. For example, if the marker genes of a newly reported cell type are co-selectively-expressed in our cell clusters, this new cell type and marker genes are added to the cell type dictionary and then included in the downstream seed cell identification and CellRef construction.
iii. *Seed cell identification*. In this step, we aim to identify cells that best represent the identity of each cell type using “single cell ranking” based on marker genes in the dictionary. These cells will then serve as “seeds” to construct the CellRef. For a cell type *i*, we first identify cells with expression of any negative markers of *i* or expressed less than two positive markers of *i* and remove those cells from *A*_*i*_ (the candidate cell clusters of cell type *i* that we identified in step 2). Using the remaining cells in *A*_*i*_, we perform “single cell ranking” using the positive markers of *i* as described in step 2 and generate an aggregated ranking of cells. Top-ranked cells in the aggregated list will be selected as the “seed” cells for cell type *i*.
iv. *Consensus prediction*. Once all “seed” cells are identified, we use them to predict cell type annotations of all cells in the collection using two independent automated cell type annotation algorithms, Seurat’s label transfer^5^ and SingleR^6^. For the Seurat’s “label transfer” based prediction, we integrate scRNA-seq data of the “seed” cells using SCTransform normalization based reciprocal principal component analysis (RPCA) integration, perform SCTransform normalization on gene expression in each of our collected datasets, and predict cell type annotations using the “MapQuery” function in Seurat v4. A predicted cell type and an associated prediction score were assigned to each query cell based on transcriptomic similarity between the query cell and the “seed” cells. Cells with low prediction scores were excluded from the CellRef construction. For the SingleR based prediction, we normalize gene expression in the “seed” cells and in a query dataset by total UMIs per cell and use the “SingleR” function with default parameters to predict cell type annotations for the query cells. We removed poor-quality or ambiguous predictions using the “pruneScores” function. Let *Y* be the set of cells with consistent cell type predictions in both methods. We calculated a *k*-nearest-neighbor purity (kNN-purity) metric for each cell in *Y*, measuring the percentage of the cell’s *k* nearest neighbors (by default, *k*=20) that have the same cell type prediction. The CellRef was comprised of the “seed” cells and the cells that have consistent cell type predictions in both methods and with kNN-purity great than 0.6.

### Construction of the “LungMAP Human Lung CellRef”

We constructed a cell type dictionary for normal human lung (a list of cell types and their associated marker genes) based on the cell types and marker genes listed in the LungMAP CellCards^10^. In addition, we extended the dictionary to include seven human lung cell types reported in recent single cell studies but not yet in CellCards, including systemic venous endothelial cell (SVEC), deuterosomal cell, submucosal gland (SMG) duct cell, megakaryocyte/platelets, suprabasal cell, mature dendritic cell (maDC), and respiratory airway secretory cell (RAS). In total, 48 cell types are defined in the dictionary.

Using this cell type dictionary, we performed the guided CellRef construction described above using seven scRNA-seq datasets. The original data were aligned to three versions of 10x Cell Ranger hg38 reference genome. To reduce the impact of reference genome differences on the data integration, we used the expression of 32,278 common gene features (based on Ensembl IDs) among the three reference genome versions to perform data integration and candidate cell cluster identification as described above. A curation was performed on the candidate cell cluster assignment by inspection of marker genes expression in the cell clusters. Based on the curated candidate cell clusters for each cell type, we selected up to top 200 cells with the lowest scores as the “seed” cells for a cell type. In total, 8,080 “seed” cells were identified for 48 normal human lung cell types. We named this collection of “seed” cells as “LungMAP Human Lung CellRef Seed”. To facilitate the use of the “LungMAP Human Lung CellRef Seed” for automated cell type annotation, we normalized gene expression in “seed” cells of each datasets using SCTransform, integrated data from different datasets using the RPCA pipeline, and performed UMAP analysis on the integrated data.

We performed a power analysis and determined the minimum cell numbers required for a lung cell type to achieve a power>=0.8. The analysis was performed as follows. First, a Cohen’s *d* effect size was calculated for each cell type using the averaged mean expression and variance of all genes in the cell type of each individual donor when compared to those in all the other cells. We grouped effect size values to the following categories: small (0.2 ≤ *d* < 0.5), medium (0.5 ≤ *d* < 0.7), large (*d* ≥ 0.7) and then used the gPower software to calculate a sample size required by each cell type using the following parameters: alpha=0.01, two tailed *t* test, beta=0.2, allocation ration=1. Based on the calculation, a minimum of 50 cells is required to reach the statistical power. 44 out of the 48 human lung cell types meet the criterial; 4 cell types had less than 50 “seed” cells identified, including chondrocytes (n=6), ILC (n=14), megakaryocyte/platelets (n=29), maDC (n=34).

Using the identified “seed” cells, we further predicted cell type annotations for all other cells in the 10 datasets collected. Both Seurat’s label transfer and SingleR were applied as described above. The “LungMAP Human Lung CellRef” (n=347,970 cells) was comprised of the “seed” cells and the cells with consistent cell type predictions and with “kNN-purity” scores >=0.6. 157,286 cells that did not pass the criteria were not included. To facilitate the use of the “LungMAP Human Lung CellRef” for automated cell type annotation, we normalized gene expression in each donor in the CellRef using SCTransform, integrated data from different donors using the RPCA pipeline, and performed UMAP analysis on the integrated data. During the RPCA integration, we identified “anchors” using the “FindIntegrationAnchors” function, filtered out “anchors” mapping cells with different cell type predictions, and then used the remaining “anchors” for data integration using the “IntegrateData” function.

### Construction of the “LungMAP Mouse Lung Development CellRef”

We constructed a cell type dictionary for mouse lung based on our constructed dictionary derived from the LungMAP CellCards. In addition, because of the developmental design of the mouse data, we extended the mouse lung cell type dictionary to include progenitor and transitional cells reported in recent single cell studies, including *Sox9*+/*Id2*+ distal epithelial cells^23,24^, “AT1/AT2” cell, *Foxf1*+/*Kit*+ endothelial progenitor cells^29^, and proliferative mesenchymal progenitor cells^30,31^. We used Seurat to perform SCTransform based data normalization and performed UMAP analysis on the identified “LungMAP Mouse Lung Development CellRef Seed” and the constructed “LungMAP Mouse Lung Development CellRef”.

### CellRef evaluation of automated cell type annotations

We developed an R script to evaluate cell type annotations predicted using the LungMAP CellRefs. Currently, the functions include: (i) Dotplot visualization of expression levels and frequencies of CellRef marker genes in each of the predicted cell types. Selective and abundant expression of marker genes in their corresponding cell types indicate a concordance of cell identities in the predictions and in the CellRef. (ii) Identification of signature genes for each of the predicted cell types. By default, the identifications were performed using Seurat’s FindAllMarkers function based on the following criteria: adjusted p value of Wilcoxon rank sum test <0.1, pct>=20%, and fold change>=1.5. A sufficient number of signature genes (e.g., >=50 genes) would be expected to define a distinct cell type. (iii) Gene sets functional enrichment analysis (Gene Ontology Biological Process, Pathways) associated with the identified cell type signature genes. Functional enrichment analysis was performed using R package gprofiler2. Given an scRNA-seq data with automated cell type annotations, the R script can generate the visualizations and evaluations for all predicted cell types at once and compile results into an evaluation report using R markdown.

## ACKNOWLEDGMENTS

We thank all members of the LungMAP2 consortium for their input and discussion. We thank the LungMAP2 external advisory committee for their insights. We thank Scott Randell and Marsico Lung Institute Tissue Procurement and Cell Culture Core for providing the human bronchial samples for the submucosal gland scRNA-seq data generation.

## AUTHOR CONTRIBUTIONS

Conceptualization: M.G. and Y.X. Writing: M.G, Y.X. and J.A.W. Data collection and processing: M.G., M.P.M, Y.W., S.Z., A.W. New data generation: J.A.W., K.A.W-B., J.A.K., K.M., A.P.N., E.E.M., K.S., M.C.B., S.M.L., Y.Y. CellRef construction: M.G. and Y.X. Web portals: Y.D., M.P.M., A.W., M.K., K.J., N.G., A.B., B.J.A., T.L.T, N.S. Cell identity and markers selection: M.G., Y.X., X.S., E.E.M., J.A.W., B.J.A., G.S.P., R.S.M., J.N.A., G.C. Project Administration: S.L. Funding Acquisition: J.A.W., Y.X., X.S., E.E.M, G.S.P., J.N.A., G.C., B.J.A., T.L.T., and N.S. All authors reviewed and approved the final version.

## DATA AVAILABILITY

The LungMAP Human Lung CellRef was constructed using data from eight published single cell/nuclei RNA-seq datasets and two unpublished single cell RNA-seq datasets from LungMAP consortium. The published datasets are accessible on Gene Expression Omnibus (access numbers: GSE135893, GSE136831, GSE122960, and GSE134174, GSE161382, GSE171524), European Genome-phenome Archive (accession number: EGAS00001004082), and Synapse.org (accession number: syn21041850). Raw sequencing data of LungMAP mouse lung CellRef is available through LungMAP.net. Processed data are accessible through LungMAP.net and LGEA web portal https://research.cchmc.org/pbge/lunggens/CellRef/LungMapCellRef.html.

## CODE AVAILABILITY

The code of LungMAP CellRef construction pipeline is available on GitHub (https://github.com/xu-lab/CellRef). The code of scViewer-lite is https://github.com/Morriseylab/scViewer-Lite.

