## Supplementary Figures 1-5 for "Guided construction of single cell reference for human and mouse lung"

Guo et al.

Supplementary Figures

**Supplementary Figure 1. Data integration and identification of the “LungMAP Human Lung CellRef Seed”.** (**A**) Integration of cells from individual donors of seven scRNA-seq datasets. (**B**) Cells colored by regions of tissue samples. (**C**) Cell clusters identified using the Leiden algorithm. (**D**) The identified “seed” cells for each of the 48 cell types in the dictionary.

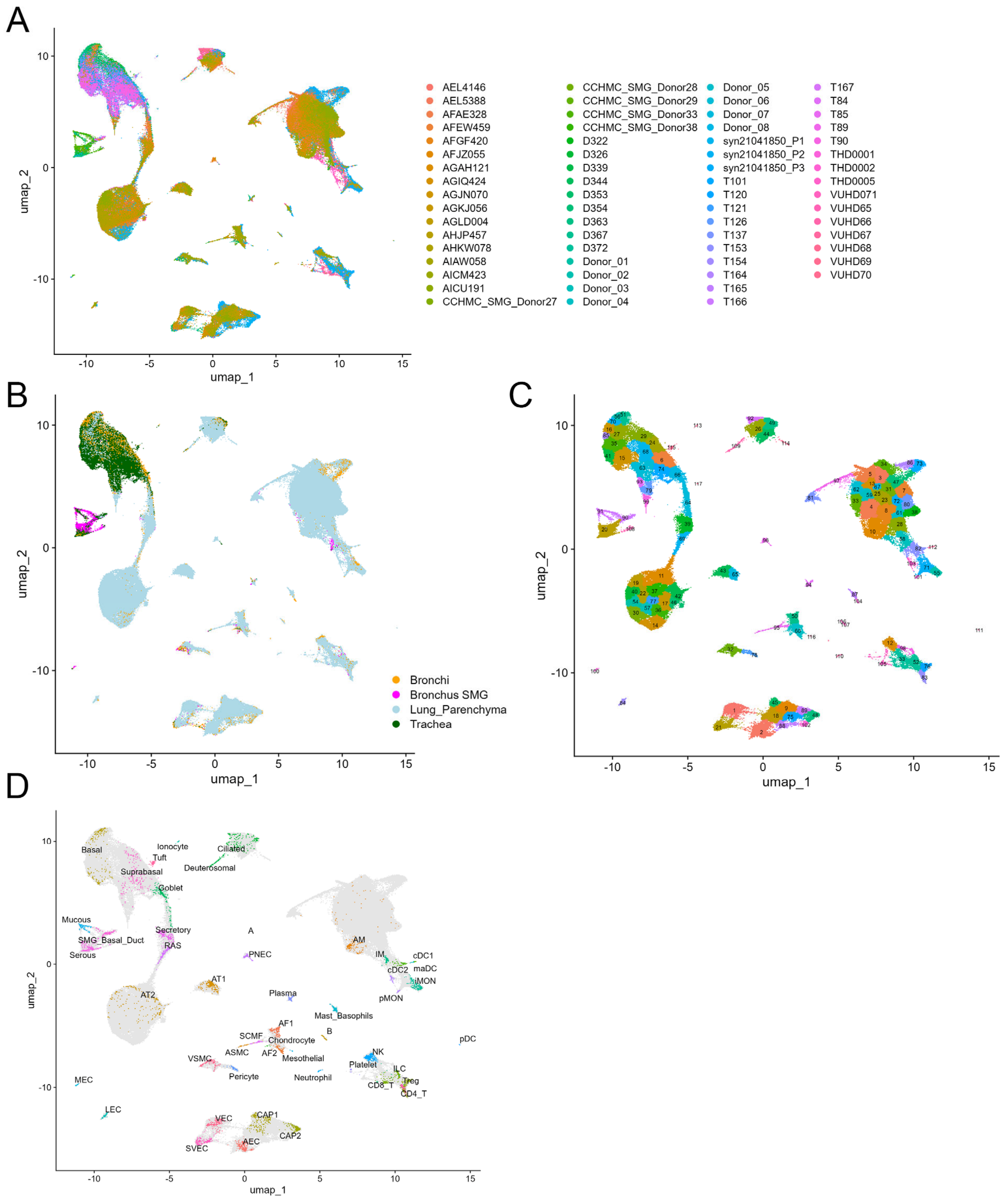

**Supplementary Figure 2. Cellular composition and cell type marker gene expression in the constructed "LungMAP Human Lung CellRef".** (**A**) Dotplot visualization of expression of cell type marker genes from the LungMAP CellCards in each cell type of the "LungMAP Human Lung CellRef". Gene expression was measured by unique molecular identifier (UMI) and normalized using Seurat's LogNormalize function. (**B**) The number of cells in each cell type in the "LungMAP Human Lung CellRef".

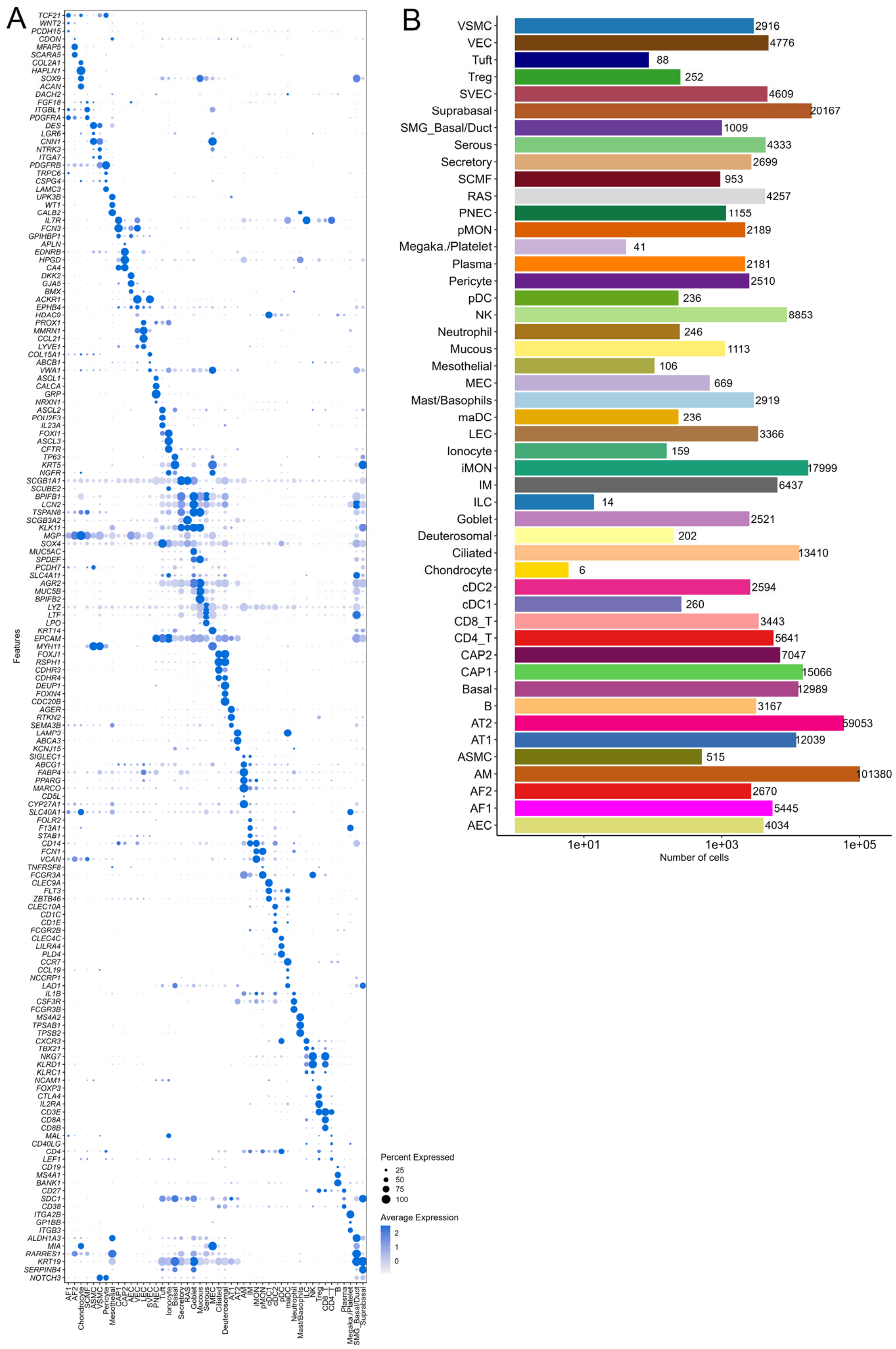

**Supplementary Figure 3. Data integration and identification of the "LungMAP Mouse Lung Development CellRef Seed".** (**A**) UMAP visualization of integrated data from 17 Drop-seq samples from eight time points of mouse lung development. (**B**) Cells colored by developmental time points. (**C**) Cell clusters identified using the Leiden algorithm. (**D**) The identified "seed" cells for each of the 40 cell types in developmental mouse lung.

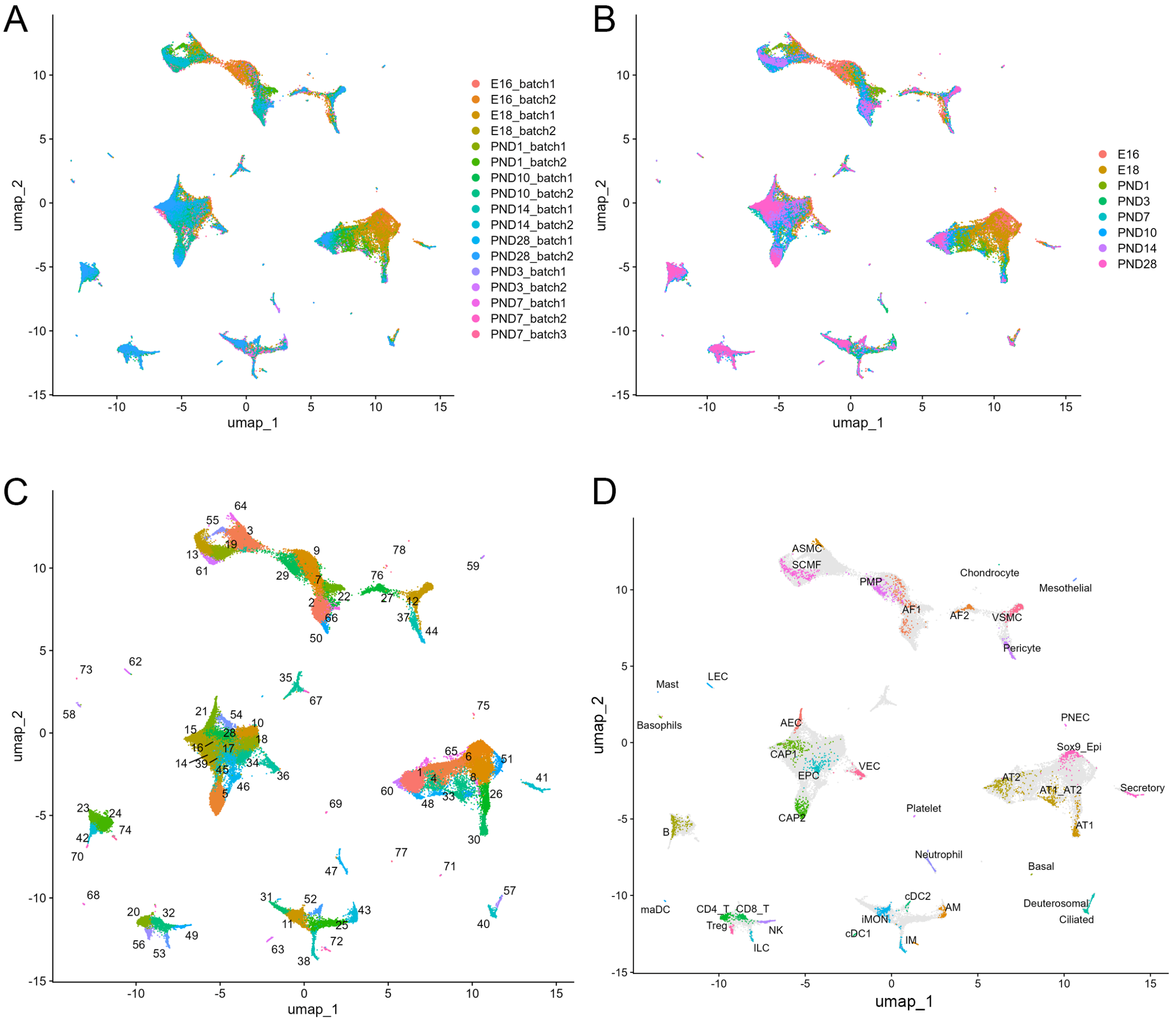

**Supplementary Figure 4. Interactive exploration of LungMAP CellRefs using scViewer-lite.** (**A**) Using scView-lite to query for the expression pattern of a single gene. Left panel shows user interfaces to enter the query. Right panels show UMAP plots of the expression of the query gene and the cell types in the "LungMAP Human Lung CellRef Seed". (**B**) Side-by-side UMAP visualizations of expression of two genes of interests using scViewer-lite.

**A.**

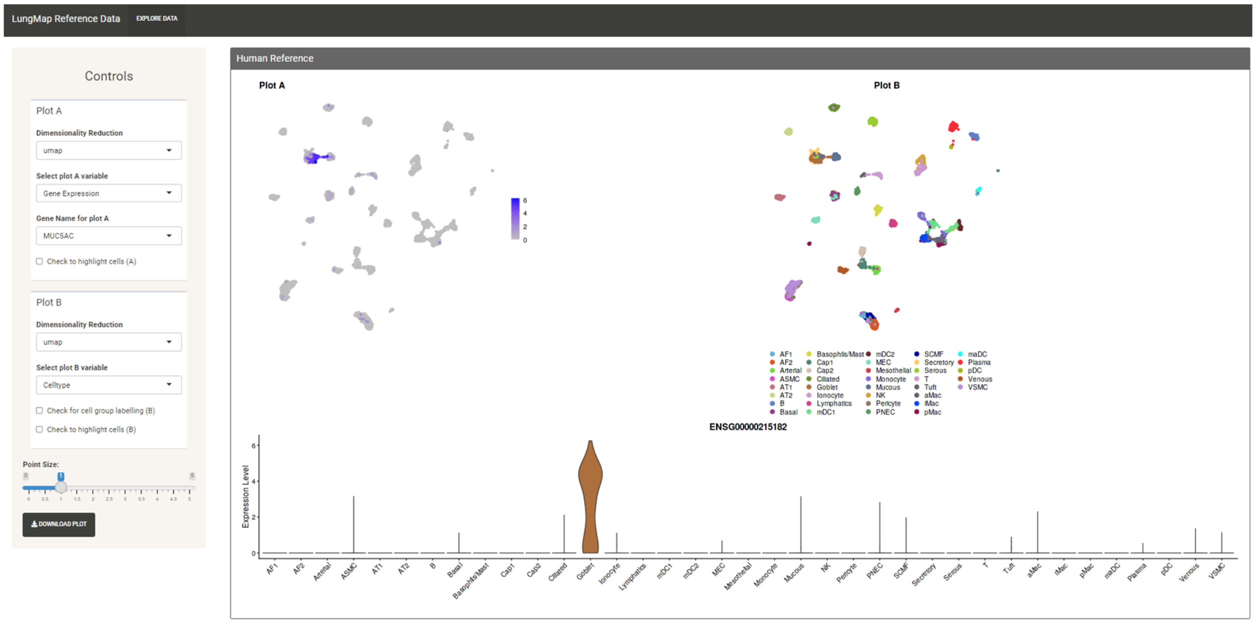


**B.**

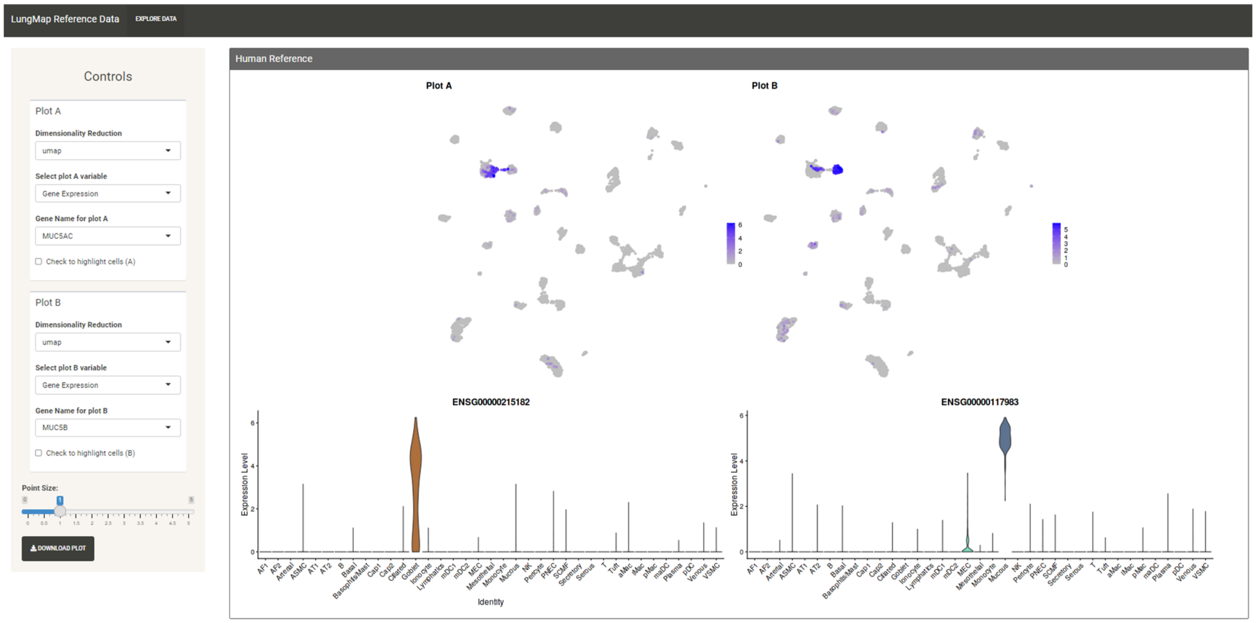

**Supplementary Figure 5. Evaluation of automated cell type annotations for scRNA-seq of normal human lung using the LungMAP CellRefs. (A)** UMAP of cells from scRNA-seq of a 2-month-old normal human lung (GSM4504966 and GSM4504967). (**B**) Dotplot visualization of expression of CellRef marker genes in the predicted cell types in A. (**C**) UMAP of cells from scRNA-seq of a 31-year-old normal human lung (GSM4035472). (**D**) Dotplot visualization of expression of CellRef marker genes in the predicted cell types in C. In (A) and (C), cells were colored by cell type annotations predicted using the “LungMAP Human Lung CellRef Seed” as reference; visualizations showed cells with prediction scores>=0.6 and predicted cell types with at least 5 cells.

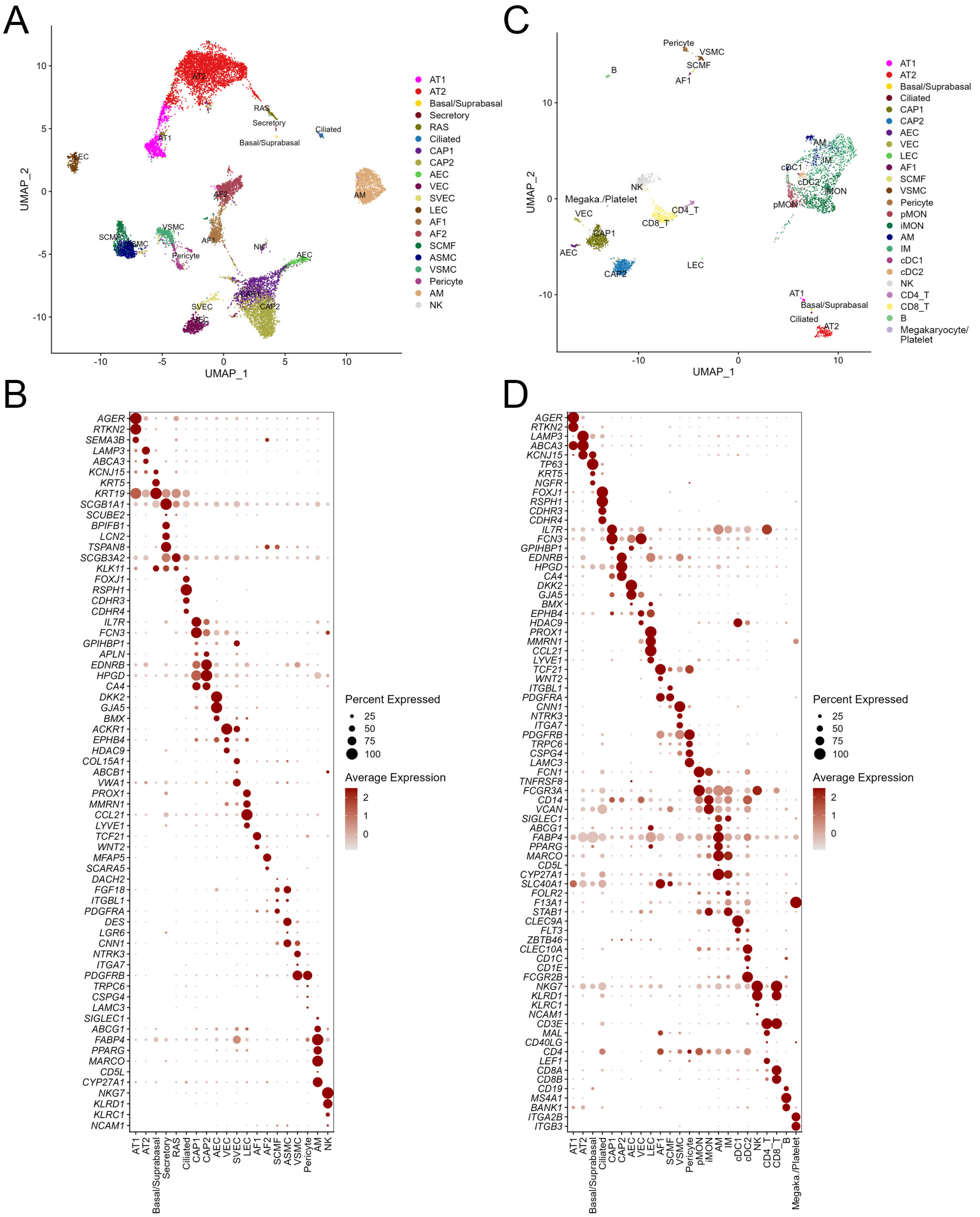
