## Supplementary Table 1 for "Guided construction of single cell reference for human and mouse lung"

**Supplementary Table 1. Collection of sc/snRNA-seq of human lung**

| Dataset | DonorID | Sex | Age |
|---|---|---|---|
| CCHMC_LungMAP | CCHMC_SMG_Donor27 | M | 47 |
| CCHMC_LungMAP | CCHMC_SMG_Donor28 | M | 20 |
| CCHMC_LungMAP | CCHMC_SMG_Donor29 | F | 13 |
| CCHMC_LungMAP | CCHMC_SMG_Donor33 | M | 50 |
| CCHMC_LungMAP | CCHMC_SMG_Donor38 | F | 39 |
| EGAS00001004082 | D322 | F | 30 |
| EGAS00001004082 | D326 | M | 31 |
| EGAS00001004082 | D339 | M | 29 |
| EGAS00001004082 | D344 | F | 26 |
| EGAS00001004082 | D353 | F | 30 |
| EGAS00001004082 | D354 | M | 33 |
| EGAS00001004082 | D363 | F | 24 |
| EGAS00001004082 | D367 | M | 27 |
| EGAS00001004082 | D372 | F | 35 |
| GSE122960 | Donor_01 | F | 63 |
| GSE122960 | Donor_02 | M | 55 |
| GSE122960 | Donor_03 | F | 29 |
| GSE122960 | Donor_04 | F | 57 |
| GSE122960 | Donor_05 | F | 49 |
| GSE122960 | Donor_06 | F | 22 |
| GSE122960 | Donor_07 | F | 47 |
| GSE122960 | Donor_08 | M | 21 |
| GSE134174 | T101 | M | 55 |
| GSE134174 | T120 | F | 57 |
| GSE134174 | T121 | F | 23 |
| GSE134174 | T126 | F | 35 |
| GSE134174 | T137 | M | 27 |
| GSE134174 | T153 | M | 38 |
| GSE134174 | T154 | M | 61 |
| GSE134174 | T164 | M | 66 |
| GSE134174 | T165 | M | 64 |
| GSE134174 | T166 | F | 68 |
| GSE134174 | T167 | F | 66 |
| GSE134174 | T84 | M | 22 |
| GSE134174 | T85 | M | 59 |
| GSE134174 | T89 | F | 10 |
| GSE134174 | T90 | M | 44 |
| GSE135893 | VUHD69 | F | 31 |
| GSE135893 | THD0001 | M | 36 |
| GSE135893 | THD0002 | | 40 |
| GSE135893 | THD0005 | M | |
| GSE135893 | VUHD071 | M | 38 |
| GSE135893 | VUHD65 | M | 17 |
| GSE135893 | VUHD66 | M | 30 |
| GSE135893 | VUHD67 | F | 30 |
| GSE135893 | VUHD68 | M | 41 |

| | | | |
|---|---|---|---|
| GSE135893 | VUHD70 | M | 54 |
| GSE136831 | 001C | M | 22 |
| GSE136831 | 002C | F | 25 |
| GSE136831 | 003C | F | 67 |
| GSE136831 | 222C | M | 65 |
| GSE136831 | 034C | M | 49 |
| GSE136831 | 065C | F | 66 |
| GSE136831 | 081C | M | 20 |
| GSE136831 | 084C | M | 46 |
| GSE136831 | 092C | M | 29 |
| GSE136831 | 098C | F | 41 |
| GSE136831 | 133C | F | 32 |
| GSE136831 | 137C | M | 54 |
| GSE136831 | 1372C | F | 21 |
| GSE136831 | 160C | M | 64 |
| GSE136831 | 192C | F | 62 |
| GSE136831 | 208C | M | 23 |
| GSE136831 | 218C | M | 29 |
| GSE136831 | 226C | M | 32 |
| GSE136831 | 244C | M | 50 |
| GSE136831 | 253C | F | 66 |
| GSE136831 | 296C | F | 80 |
| GSE136831 | 388C | M | 61 |
| GSE136831 | 396C | F | 37 |
| GSE136831 | 439C | F | 66 |
| GSE136831 | 454C | F | 48 |
| GSE136831 | 465C | M | 56 |
| GSE136831 | 483C | M | 35 |
| GSE136831 | 484C | M | 31 |
| UPenn_LungMAP | AEL4146 | M | 52 |
| UPenn_LungMAP | AEL5388 | M | 69 |
| UPenn_LungMAP | AFAE328 | F | 35 |
| UPenn_LungMAP | AFEW459 | F | 76 |
| UPenn_LungMAP | AFGF420 | F | 51 |
| UPenn_LungMAP | AFJZ055 | M | 23 |
| UPenn_LungMAP | AGAH121 | F | 70 |
| UPenn_LungMAP | AGIQ424 | F | 29 |
| UPenn_LungMAP | AGJN070 | M | 25 |
| UPenn_LungMAP | AGKJ056 | M | 17 |
| UPenn_LungMAP | AGLD004 | M | 13 |
| UPenn_LungMAP | AHJP457 | F | 57 |
| UPenn_LungMAP | AHKW078 | F | 68 |
| UPenn_LungMAP | AIAW058 | F | 59 |
| UPenn_LungMAP | AICM423 | F | 12 |
| UPenn_LungMAP | AICU191 | M | 42 |
| syn21041850 | syn21041850_P1 | M | 75 |
| syn21041850 | syn21041850_P2 | M | 46 |
| syn21041850 | syn21041850_P3 | F | 51 |
| GSE161382 | D122 | M | 31 |

| | | | |
|---|---|---|---|
| GSE161382 | D175 | F | 29 |
| GSE161382 | D231 | F | 33 |
| GSE171524 | C51ctr | F | 70 |
| GSE171524 | C52ctr | F | 69 |
| GSE171524 | C53ctr | M | 79 |
| GSE171524 | C54ctr | F | 72 |
| GSE171524 | C55ctr | M | 69 |
| GSE171524 | C56ctr | M | 75 |
| GSE171524 | C57ctr | M | 68 |
