## Supplementary Table 2 for "Guided construction of single cell reference for human and mouse lung"

**Supplementary Table 2. Cell type dictionary for the LungMAP Human Lung CellRef construction**

| Cell type | Marker | MarkerType |
|---|---|---|
| AF1 | *TCF21* | p |
| AF1 | *WNT2* | p |
| AF1 | *PCDH15* | p |
| AF2 | *CDON* | p |
| AF2 | *MFAP5* | p |
| AF2 | *SCARA5* | p |
| Chondrocyte | *COL2A1* | p |
| Chondrocyte | *HAPLN1* | p |
| Chondrocyte | *SOX9* | p |
| Chondrocyte | *ACAN* | p |
| SCMF | *DACH2* | p |
| SCMF | *FGF18* | p |
| SCMF | *ITGBL1* | p |
| SCMF | *PDGFRA* | p |
| ASMC | *DES* | p |
| ASMC | *LGR6* | p |
| VSMC | *CNN1* | p |
| VSMC | *NTRK3* | p |
| VSMC | *ITGA7* | p |
| Pericyte | *PDGFRB* | p |
| Pericyte | *TRPC6* | p |
| Pericyte | *CSPG4* | p |
| Pericyte | *LAMC3* | p |
| Mesothelial | *UPK3B* | p |
| Mesothelial | *WT1* | p |
| Mesothelial | *CALB2* | p |
| CAP1 | *IL7R* | p |
| CAP1 | *FCN3* | p |
| CAP1 | *GPIHBP1* | p |
| CAP2 | *APLN* | p |
| CAP2 | *EDNRB* | p |
| CAP2 | *HPGD* | p |
| CAP2 | *CA4* | p |
| AEC | *DKK2* | p |
| AEC | *GJA5* | p |
| AEC | *BMX* | p |
| VEC | *ACKR1* | p |
| VEC | *EPHB4* | p |
| VEC | *HDAC9* | p |
| LEC | *PROX1* | p |
| LEC | *MMRN1* | p |
| LEC | *CCL21* | p |
| LEC | *LYVE1* | p |
| SVEC | *ACKR1* | p |
| SVEC | *COL15A1* | p |

| | | |
|---|---|---|
| SVEC | *ABCB1* | p |
| SVEC | *VWA1* | p |
| PNEC | *ASCL1* | p |
| PNEC | *CALCA* | p |
| PNEC | *GRP* | p |
| PNEC | *NRXN1* | p |
| Tuft | *ASCL2* | p |
| Tuft | *POU2F3* | p |
| Tuft | *IL23A* | p |
| Ionocyte | *FOXI1* | p |
| Ionocyte | *ASCL3* | p |
| Ionocyte | *CFTR* | p |
| Basal | *TP63* | p |
| Basal | *KRT5* | p |
| Basal | *NGFR* | p |
| Secretory | *SCGB1A1* | p |
| Secretory | *SCUBE2* | p |
| Secretory | *BPIFB1* | p |
| Secretory | *LCN2* | p |
| Secretory | *TSPAN8* | p |
| RAS | *SCGB3A2* | p |
| RAS | *KLK11* | p |
| RAS | *MGP* | p |
| RAS | *SOX4* | p |
| Goblet | *MUC5AC* | p |
| Goblet | *SPDEF* | p |
| Goblet | *PCDH7* | p |
| Goblet | *SLC4A11* | p |
| Goblet | *AGR2* | p |
| Mucous | *MUC5B* | p |
| Mucous | *SPDEF* | p |
| Mucous | *BPIFB2* | p |
| Mucous | *MUC5AC* | n |
| Serous | *LYZ* | p |
| Serous | *LTF* | p |
| Serous | *LPO* | p |
| MEC | *KRT14* | p |
| MEC | *EPCAM* | p |
| MEC | *MYH11* | p |
| Ciliated | *FOXJ1* | p |
| Ciliated | *RSPH1* | p |
| Ciliated | *CDHR3* | p |
| Ciliated | *CDHR4* | p |
| Deuterosomal | *DEUP1* | p |
| Deuterosomal | *FOXN4* | p |
| Deuterosomal | *CDC20B* | p |
| SMG Basal/Duct | *KRT14* | p |
| SMG Basal/Duct | *SOX9* | p |
| SMG Basal/Duct | *ALDH1A3* | p |

| | | |
|---|---|---|
| SMG Basal/Duct | *MIA* | p |
| SMG Basal/Duct | *RARRES1* | p |
| Suprabasal | *KRT19* | p |
| Suprabasal | *SERPINB4* | p |
| Suprabasal | *NOTCH3* | p |
| AT1 | *AGER* | p |
| AT1 | *RTKN2* | p |
| AT1 | *SEMA3B* | p |
| AT2 | *LAMP3* | p |
| AT2 | *ABCA3* | p |
| AT2 | *KCNJ15* | p |
| AM | *SIGLEC1* | p |
| AM | *ABCG1* | p |
| AM | *FABP4* | p |
| AM | *PPARG* | p |
| AM | *MARCO* | p |
| AM | *CD5L* | p |
| AM | *CYP27A1* | p |
| IM | *SLC40A1* | p |
| IM | *FOLR2* | p |
| IM | *F13A1* | p |
| IM | *STAB1* | p |
| iMON | *CD14* | p |
| iMON | *FCN1* | p |
| iMON | *VCAN* | p |
| iMON | *FCGR3A* | n |
| pMON | *FCN1* | p |
| pMON | *TNFRSF8* | p |
| pMON | *FCGR3A* | p |
| cDC1 | *CLEC9A* | p |
| cDC1 | *FLT3* | p |
| cDC1 | *ZBTB46* | p |
| cDC2 | *CLEC10A* | p |
| cDC2 | *CD1C* | p |
| cDC2 | *CD1E* | p |
| cDC2 | *FCGR2B* | p |
| pDC | *CLEC4C* | p |
| pDC | *LILRA4* | p |
| pDC | *PLD4* | p |
| maDC | *CCR7* | p |
| maDC | *CCL19* | p |
| maDC | *NCCRP1* | p |
| maDC | *LAD1* | p |
| Neutrophil | *IL1B* | p |
| Neutrophil | *CSF3R* | p |
| Neutrophil | *FCGR3B* | p |
| Mast/Basophil | *MS4A2* | p |
| Mast/Basophil | *TPSAB1* | p |
| Mast/Basophil | *TPSB2* | p |

| | | |
|---|---|---|
| ILC | *IL7R* | p |
| ILC | *CXCR3* | p |
| ILC | *TBX21* | p |
| ILC | *EOMES* | n |
| ILC | *CD3E* | n |
| ILC | *CD3D* | n |
| ILC | *CD3G* | n |
| ILC | *CD14* | n |
| ILC | *FCGR3A* | n |
| ILC | *CD19* | n |
| ILC | *MS4A1* | n |
| ILC | *NCAM1* | n |
| NK | *NKG7* | p |
| NK | *KLRD1* | p |
| NK | *KLRC1* | p |
| NK | *NCAM1* | p |
| Treg | *FOXP3* | p |
| Treg | *CTLA4* | p |
| Treg | *IL2RA* | p |
| CD8_T | *CD3E* | p |
| CD8_T | *CD8A* | p |
| CD8_T | *CD8B* | p |
| CD4_T | *CD3E* | p |
| CD4_T | *MAL* | p |
| CD4_T | *CD40LG* | p |
| CD4_T | *CD4* | p |
| CD4_T | *LEF1* | p |
| B | *CD19* | p |
| B | *MS4A1* | p |
| B | *BANK1* | p |
| Plasma | *CD27* | p |
| Plasma | *SDC1* | p |
| Plasma | *CD38* | p |
| Megakaryocyte/Platelet | *ITGA2B* | p |
| Megakaryocyte/Platelet | *GP1BB* | p |
| Megakaryocyte/Platelet | *ITGB3* | p |
