## Supplementary Table 3 for "Guided construction of single cell reference for human and mouse lung"

**Supplementary Table 3. Drop-seq of Mouse Lung data**

| Time | Sample | # Cells |
|---|---|---|
| E16.5 | E16_batch1 | 9051 |
| E16.5 | E16_batch2 | 13428 |
| E18.5 | E18_batch1 | 7630 |
| E18.5 | E18_batch2 | 5138 |
| PND1 | PND1_batch1 | 2307 |
| PND1 | PND1_batch2 | 5271 |
| PND3 | PND3_batch1 | 7077 |
| PND3 | PND3_batch2 | 3406 |
| PND7 | PND7_batch1 | 1090 |
| PND7 | PND7_batch2 | 5477 |
| PND7 | PND7_batch3 | 5934 |
| PND10 | PND10_batch1 | 6653 |
| PND10 | PND10_batch2 | 7257 |
| PND14 | PND14_batch1 | 2964 |
| PND14 | PND14_batch2 | 5654 |
| PND28 | PND28_batch1 | 2420 |
| PND28 | PND28_batch2 | 4901 |
