## Supplementary Table 4 for "Guided construction of single cell reference for human and mouse lung"

**Supplementary Table 4. Cell type dictionary of the LungMAP Mouse Lung Development CellRef construction**

| Cell type | Marker | MarkerType |
|---|---|---|
| AF1 | *Tcf21* | p |
| AF1 | *Wnt2* | p |
| AF1 | *Mfap4* | p |
| AF2 | *Cdon* | p |
| AF2 | *Mfap5* | p |
| AF2 | *Scara5* | p |
| AF2 | *Col14a1* | p |
| Chondrocyte | *Col2a1* | p |
| Chondrocyte | *Hapln1* | p |
| Chondrocyte | *Sox9* | p |
| Chondrocyte | *Acan* | p |
| SCMF | *Fgf18* | p |
| SCMF | *Pdgfra* | p |
| SCMF | *Acta2* | p |
| ASMC | *Acta2* | p |
| ASMC | *Des* | p |
| ASMC | *Lgr6* | p |
| ASMC | *Hhip* | p |
| VSMC | *Cnn1* | p |
| VSMC | *Ntrk3* | p |
| VSMC | *Itga7* | p |
| Pericyte | *Pdgfrb* | p |
| Pericyte | *Trpc6* | p |
| Pericyte | *Cspg4* | p |
| Mesothelial | *Upk3b* | p |
| Mesothelial | *Wt1* | p |
| Mesothelial | *Frem2* | p |
| Mesothelial | *Lrrn4* | p |
| CAP1 | *Aplnr* | p |
| CAP1 | *Gpihbp1* | p |
| CAP1 | *Ptprb* | p |
| CAP2 | *Apln* | p |
| CAP2 | *Ednrb* | p |
| CAP2 | *Hpgd* | p |
| CAP2 | *Car4* | p |
| AEC | *Dkk2* | p |
| AEC | *Gja5* | p |
| AEC | *Bmx* | p |
| VEC | *Ephb4* | p |
| VEC | *Slc6a2* | p |
| VEC | *Ackr3* | p |
| LEC | *Prox1* | p |
| LEC | *Mmrn1* | p |
| LEC | *Ccl21a* | p |

| | | |
|---|---|---|
| LEC | *Nrp2* | p |
| PNEC | *Ascl1* | p |
| PNEC | *Calca* | p |
| PNEC | *Nrxn1* | p |
| Basal | *Trp63* | p |
| Basal | *Krt5* | p |
| Secretory | *Scgb1a1* | p |
| Secretory | *Cyp2f2* | p |
| Ciliated | *Foxj1* | p |
| Ciliated | *Rsph1* | p |
| Ciliated | *Cdhr3* | p |
| Ciliated | *Cdhr4* | p |
| Deuterosomal | *Deup1* | p |
| Deuterosomal | *Foxn4* | p |
| Deuterosomal | *Cdc20b* | p |
| AT1 | *Ager* | p |
| AT1 | *Rtkn2* | p |
| AT1 | *Sema3b* | p |
| AT2 | *Lamp3* | p |
| AT2 | *Abca3* | p |
| AT2 | *Kcnj15* | p |
| AM | *Siglec1* | p |
| AM | *Abcg1* | p |
| AM | *Fabp4* | p |
| AM | *Pparg* | p |
| AM | *Marco* | p |
| AM | *Cd5l* | p |
| AM | *Cyp27a1* | p |
| IM | *Slc40a1* | p |
| IM | *Folr2* | p |
| IM | *Ms4a6d* | p |
| IM | *F13a1* | p |
| IM | *Stab1* | p |
| iMON | *Fcnb* | p |
| iMON | *Ly6c2* | p |
| iMON | *Itgam* | p |
| cDC1 | *Clec9a* | p |
| cDC1 | *Flt3* | p |
| cDC1 | *Zbtb46* | p |
| cDC2 | *Clec10a* | p |
| cDC2 | *Fcgr2b* | p |
| maDC | *Ccr7* | p |
| maDC | *Lad1* | p |
| Neutrophil | *Il1b* | p |
| Neutrophil | *Csf3r* | p |
| Neutrophil | *Fcgr3* | p |
| Neutrophil | *Ly6g* | p |
| Mast | *Ms4a2* | p |
| Mast | *Tpsab1* | p |

| | | |
|---|---|---|
| Mast | *Tpsb2* | p |
| Mast | *Kit* | p |
| Basophil | *Cd200r3* | p |
| Basophil | *Fcer1a* | p |
| Basophil | *Itga2* | p |
| Basophil | *Kit* | n |
| ILC | *Il7r* | p |
| ILC | *Gata3* | p |
| ILC | *Thy1* | p |
| ILC | *Eomes* | n |
| ILC | *Cd3e* | n |
| ILC | *Cd3d* | n |
| ILC | *Cd3g* | n |
| ILC | *Cd14* | n |
| ILC | *Fcgr3* | n |
| ILC | *Cd19* | n |
| ILC | *Ms4a1* | n |
| ILC | *Ncam1* | n |
| NK | *Nkg7* | p |
| NK | *Klrd1* | p |
| NK | *Klrc1* | p |
| Treg | *Foxp3* | p |
| Treg | *Ctla4* | p |
| Treg | *Il2ra* | p |
| CD8_T | *Cd3e* | p |
| CD8_T | *Cd8a* | p |
| CD8_T | *Cd8b1* | p |
| CD4_T | *Cd3e* | p |
| CD4_T | *Cd40lg* | p |
| CD4_T | *Cd4* | p |
| CD4_T | *Lef1* | p |
| B | *Cd19* | p |
| B | *Ms4a1* | p |
| B | *Bank1* | p |
| Megakaryocyte/Platelet | *Itga2b* | p |
| Megakaryocyte/Platelet | *Gp1bb* | p |
| Megakaryocyte/Platelet | *Itgb3* | p |
| Sox9_Epi | *Sox9* | p |
| Sox9_Epi | *Id2* | p |
| Sox9_Epi | *Epcam* | p |
| AT1/AT2 | *Lamp3* | p |
| AT1/AT2 | *Abca3* | p |
| AT1/AT2 | *Rtkn2* | p |
| AT1/AT2 | *Sema3b* | p |
| AT1/AT2 | *Cldn4* | p |
| EPC | *Kit* | p |
| EPC | *Foxf1* | p |
| EPC | *Dll4* | p |
| EPC | *Mki67* | p |

| | | |
|---|---|---|
| PMP | *Top2a* | p |
| PMP | *Col1a1* | p |
| PMP | *Col1a2* | p |
| PMP | *Runx1t1* | p |
| PMP | *Vcan* | p |
| PMP | *Hoxb5* | p |
| PMP | *Snai2* | p |
